# Making plant tissue accessible for cryo-electron tomography

**DOI:** 10.1101/2025.02.14.638237

**Authors:** Matthias Pöge, Marcel Dickmanns, Peng Xu, Meijing Li, Oda H. Schiøtz, Christoph O. J. Kaiser, Jianfei Ma, Anna Bieber, Cristina Capitanio, Johann Brenner, Margot Riggi, Sven Klumpe, Manuel Miras, Neda S. Kazemein Jasemi, Waltraud X. Schulze, Rüdiger Simon, Wolf B. Frommer, Jürgen M. Plitzko, Wolfgang Baumeister

## Abstract

Cryo-electron tomography (cryo-ET) allows to visualize the molecular architecture of pristinely preserved cells and tissues in situ. The workflow of sample preparation for cryo-ET is rather complex; it involves vitrification by rapid freezing followed by cryo-focused ion beam (FIB) milling rendering the volumes of interest thin enough for cryo-ET data acquisition. The established protocols for single cells grown on or deposited on EM-grids are not suitable for multicellular plant tissues. Plunge-freezing does not yield vitrified samples in most cases and must be replaced by high-pressure freezing. This, in turn, necessitates extensive modifications of the subsequent FIB milling procedures. In this communication we describe procedures for sample screening, targeted FIB milling guided by cryo-fluorescence light microscopy and a novel lamella trimming step that allows to obtain homogenously thin lamellae suitable for cryo-ET. We have tested all the steps along the workflow with a variety of plant tissues including the moss *Physcomitrium patens*, *Arabidopsis thaliana* seed embryos and *Limonium bicolor* salt glands. We could demonstrate that the workflow optimized for plant tissues allows to attain subnanometer resolution in cases where subtomogram averaging is applicable.

## Main text

New imaging modalities in conjunction with advances in sample preparation can pave the way for unprecedented insights into the structural organization of cells and tissues. Cryo-electron tomography (cryo-ET), i.e. the application of tomographic principles of image acquisition and reconstruction to biological materials in a frozen-hydrated state, allows to visualize the molecular architecture of unperturbed cellular environments^1^. It combines the power of high-resolution three-dimensional (3D) imaging with close-to-life structural preservation and provides faithful representations of the physiological state of cells^2^.

Instead of disrupting cells, fractioning their molecular inventory, and studying its components in isolation, cryo-ET allows to visualize them *in situ* embedded in their functional environments. Thereby, the network of interactions underlying cellular physiology is preserved and can be revealed^3^. Moreover, many supramolecular assemblies cannot be isolated without compromising their structural integrity. Advances in technology and methodology have established structural biology in cells or *in situ* as a viable alternative^4^. Today, cryo-ET is widely deployed and increasingly popular at the interface of molecular biology and cell biology – with exception of the plant sciences, where technical issues have hampered its use.

The most critical step in cryo-ET is the vitrification. For plant tissues, this process is complicated by the cell size (diameter typically 10 – 100 µm) which is larger than for most yeast, algae or mammalian cells. Plant cells often contain water-rich vacuoles^5^ occupying up to 90 % of their cellular volume^6^ which poorly vitrify^7^ and, in the case of leaves, the tissue comprises air-filled spaces. Plunge-freezing^8^, the most convenient way to achieve vitrification, works reliably only for cells smaller than 10 µm. While cryoprotectants^9^ may be used to increase vitrification depth, for larger cells and tissues, High-pressure freezing^10^ (HPF) must be applied and still not always guarantees full vitrification. Moreover, it is challenging to make HPF compatible with the subsequent step of the cryo-ET workflow: thinning the sample to make it suitable for electron tomography data acquisition. This is commonly achieved by focused ion-beam (FIB) milling, where material is ablated from the ice-embedded sample until thin sheets (lamellae) with thicknesses of 100 – 300 nm are obtained^11^. To ensure that features of interest are included in the lamella, FIB milling is often guided by cryo-fluorescence light microscopy (FLM), pinpointing the spatial location of structures of interest. For this correlative light and electron microscopy (CLEM) approach it is possible to use FLMs equipped with cryo-compatible stages^12^ (e.g. Leica SP8/Stellaris or Linkam CMS196 stage) or a FLM integrated into the FIB chamber (int-FLM, e.g. Delmic METEOR^13^ or Thermo Fisher Scientific iFLM^14^). Recently, progress has been made in the preparation of lamellae from HPF samples for cryo-ET^15–23^. These new procedures, however, have not yet been put into practice for intact plant tissues. Previous cryo-ET work focused on Protoplasts^24^, i.e. isolated cells without cell wall, isolated organelles^25^, cyanobacteria^26^, red algae^27^ and single celled green algae^28,29^.

Structural studies of intact plant systems have opted for an alternative route, namely HPF followed by freeze substitution (HPF-FS). In this process, high-pressure frozen samples are metal-impregnated and dehydrated upon embedding in a resin allowing sectioning and imaging at room temperature^30^. Currently, this is the most accessible and reliable approach to image plant tissues by electron microscopy^7,31^. While HPF-FS generally preserves cellular ultrastructure better than chemical fixation followed by metal staining and resin embedding^32^, it is impossible to ascertain that vitrification has been adequately achieved in the first instance. To visualize and interpret structures at the molecular level, the only suitable method is cryo-ET, with samples unadulterated by dehydration and chemical modifications.

In this study, we describe workflows suitable for the preparation of plant tissues for cryo-ET. First, we applied them to protonemata and phyllids of the moss *Physcomitrium patens*^33^ forming chains and sheets, respectively, of cylindrical cells with diameters of 15-30 µm. We employed FIB-milling guided by FLM, specifically targeting cell junctions, to show that features small compared to the size of whole cells can be addressed. Later we adapted the methods to *Arabidopsis thaliana* seed embryos and *Limonium bicolor* salt glands to demonstrate the potential of the workflows, which we believe are not limited to plants but widely applicable also to other multicellular systems.

## Results

### Plunge-freezing does not vitrify *Physcomitrium patens* tissues

First, we applied the standard plunge-freezing and FIB milling workflow to *P. patens* protonemata and phyllids. As the cells were exposed on the surface of grids, the identification of cell junctions was possible using secondary electron images derived with the scanning electron microscope (SEM) and FIB, without the need for fluorescence targeting by correlation to FLM data (Figure 1 – figure supplement 4). Overviews of lamellae recorded in a transmission electron microscope (TEM), however, revealed that the cells were not vitreous (Figure 1a, N = 15). The formation of large ice crystals during plunge-freezing resulted in distortion of the cellular architecture and the generation of severe TEM imaging artefacts.

**Figure 1.**
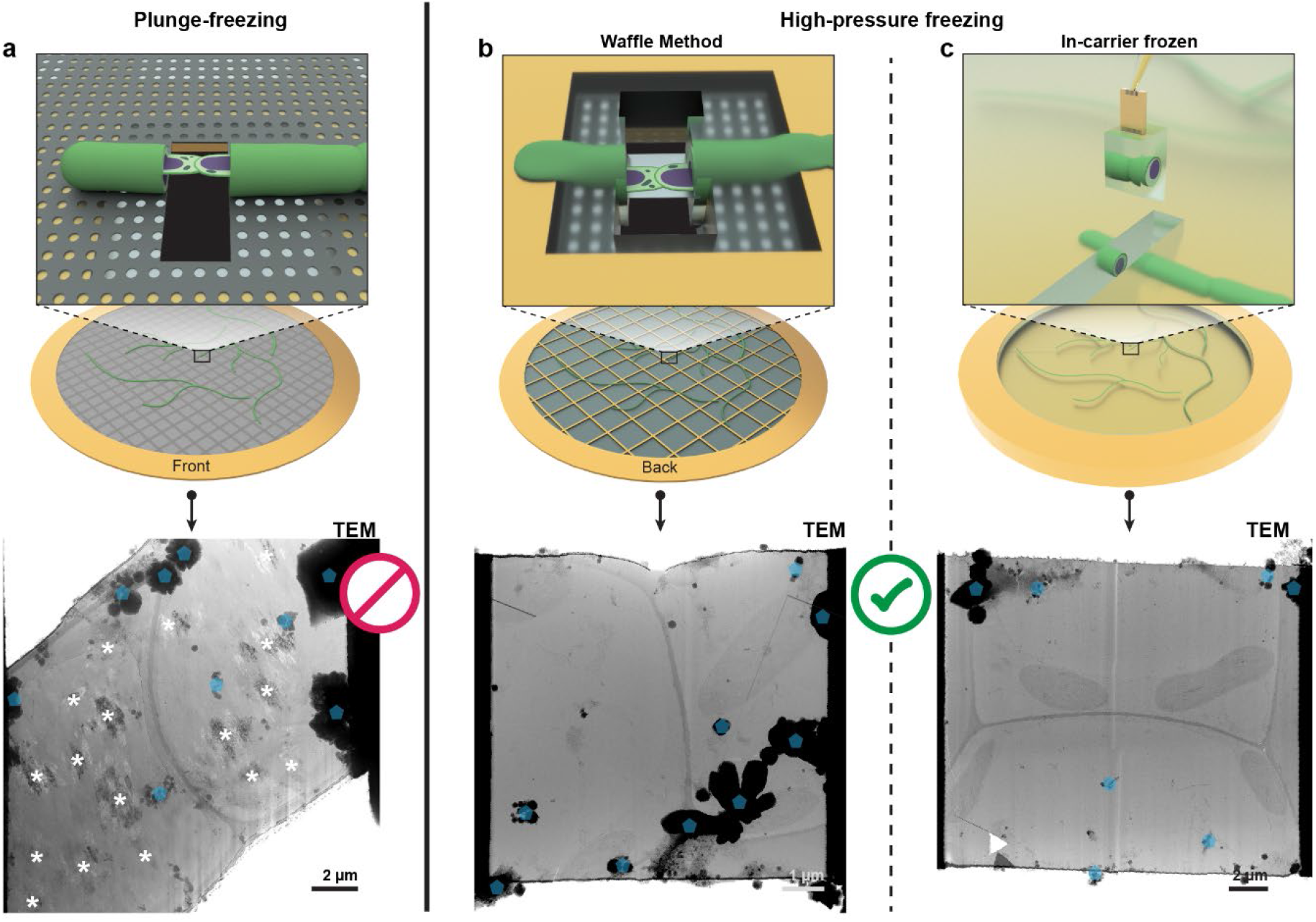
Plunge-freezing versus high-pressure freezing (HPF) for the moss *Physcomitrium patens*. The top and bottom row show schemes of the samples and corresponding examples of TEM overview images, respectively. **a**, Plunge-freezing. Patterns introduced by crystalline ice are indicated by asterisks (*). **b**,**c**, High-pressure freezing. Samples prepared by the Waffle Method or in-carrier HPF in **(b)** and **(c)**, respectively. TEM overviews of lamellae from HPF samples show no indication of crystalline ice. In (a)-(c), ice crystals contaminating the lamella surface which were introduced during transfers between microscopes are marked by blue transparent pentagons. The respective workflows are summarized in Figure 1 – figure supplement 1 and 2.

We also performed plunge-freezing with glycerol, sucrose and proline additives serving as cryoprotectants (Table 1). These, however, did either not prevent crystalline ice formation or induce plasmolysis (data not shown). In some cases, the tissue was even severely dehydrated (Figure 1 – figure supplement 4f). Thus, we concluded that it is challenging to vitrify *P. patens* tissues by plunge-freezing. While the success of plunge-freezing is sample dependent, we anticipate similar vitrification problems for vascular plants and mosses with cells of similar or larger size.

**Table 1.** Plunge-freezing of *P. patens* with cryoprotectants. List of the tested plunge-freezing methods with the employed concentrations of cryoprotectants.

|  | Culture on Solid BCDAT medium with |  | Incubation in liquid BCDAT medium with |  |  | Before plunging, add 3 µL BCDAT with |  |
| --- | --- | --- | --- | --- | --- | --- | --- |
|  | Cryoprotectant | Concentration | Cryoprot. | Conc. (w/V) | Inc. time | Cryoprotectant | Concentration |
| Method 1 |  |  |  |  |  | Sucrose | 10 %, 15 %, 20 % (w/V) |
| Method 2 | Proline | 50 mM, 75 mM, 100 mM |  |  |  | Proline | 50 mM, 75 mM, 100 mM |
| Method 3 | Proline | 50 mM, 75 mM, 100 mM | Sucrose | 10 %, 20 % | 2 h |  |  |
| Method 4 |  |  |  |  |  | Glycerol | 10 % (w/V) |

### The Waffle Method utilized for *P. patens*

Next, we tested the Waffle Method^21^ to vitrify protonemata. The tissue was placed on the back of an EM-grid, i.e. on the reverse side of a continuous support film and squeezed into the buffer-filled wells formed by the EM-grid bars and support film prior to HPF. While this approach damages cells on top of grid bars, the cells within grid squares remain intact. Hence, using EM-grids with large squares (50-100 mesh) maximizes the area of intact, vitreous tissue.

In contrast to plunge-frozen samples, the cells on waffle grids are embedded in a layer of vitreous buffer and not exposed at the sample surface (Figure 2 – figure supplement 1a). Therefore, we screened waffle grids in a cryo-FLM recording the cellular autofluorescence to identify single layers of intact protonemata tissue with cell junctions in the center of grid squares (Figure 2a,b). Furthermore, squares with multiple cell layers were ignored as only a single layer of protonemata fits into the 25 µm high well of the used grids without mechanical damage caused by compression (Figure 2 – figure supplement 1b,c).

**Figure 2.**
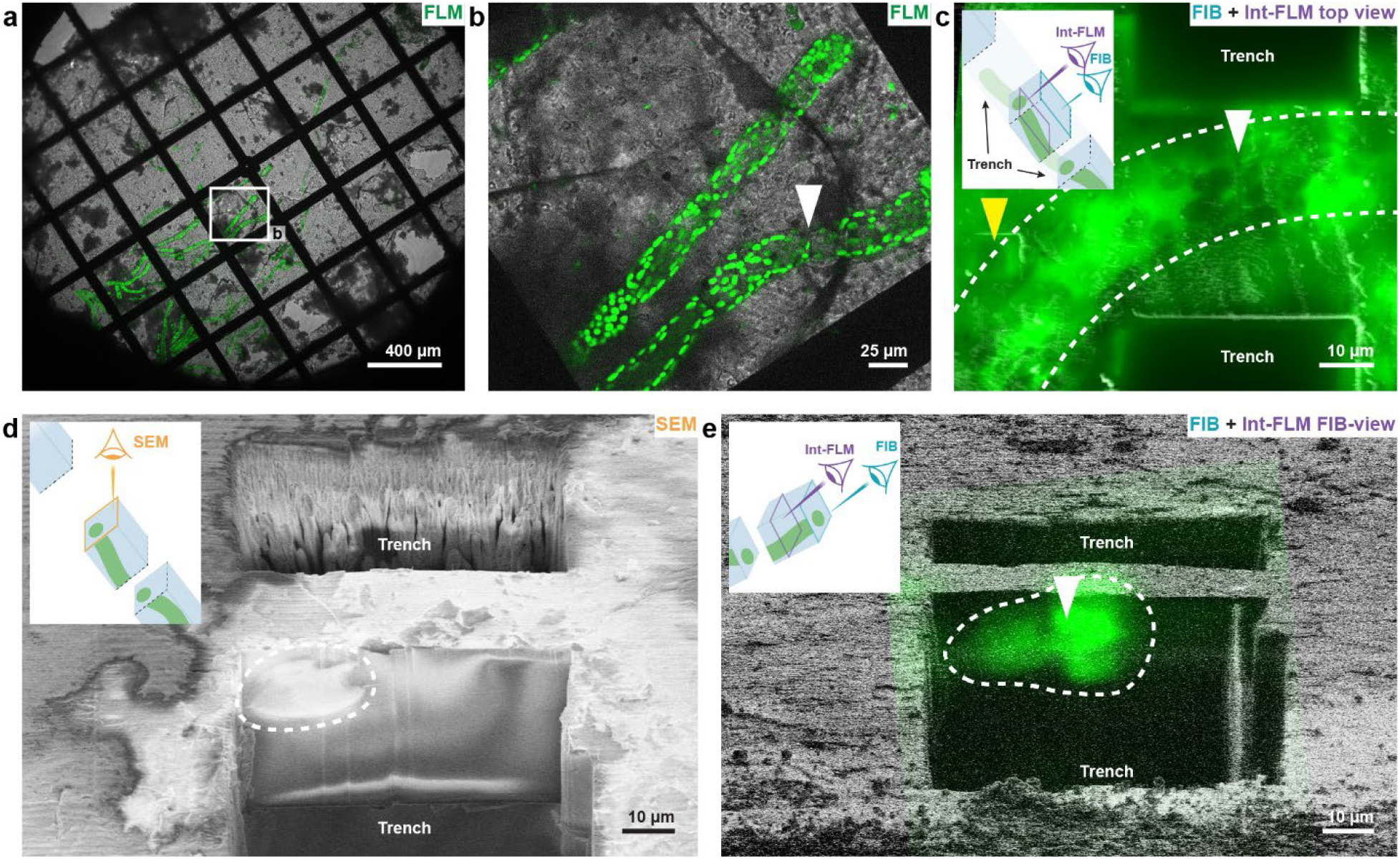
Sample screening and targeting of cell junctions in *P. patens* protonemata. **a,b**, Sample screening using a cryo-fluorescence light microscope (FLM). An overview of a waffle grid is shown in **(a)**. The white rectangle indicates the position of the grid squares in **(b)** containing a target area. Displayed in **(a)** and **(b)** are the transmitted light channel (gray) and the cellular autofluorescence (green). **c,** Lateral targeting. Overlay of a FIB image (gray) and cellular autofluorescence (green) recorded with the integrated fluorescence light microscope (int-FLM). Both images (Figure 2 – figure supplement 1g) were acquired with the stage in trench milling orientation resulting in sample top views. **d,e**, axial targeting. **d**, SEM image of the block-face exposed through trench milling. **e**, Overlay of FIB (gray) and int-FLM FIB-view image (green) of the cellular autofluorescence. Both images were recorded with the stage in lamella milling orientation. Cell junctions and locations of cells are marked with white arrowheads and dashed lines, respectively, in (**b-e**). Note that the excitation wavelengths and emission filters used for FLM imaging in **(a)** and **(b)** are different from those used for int-FLM imaging in **(c)** and **(e)**. Therefore, the fluorescence signals appear different. Ideally, for most reliable targeting, excitation and emission should be the same for all FLM imaging modalities.

Subsequently, we loaded the samples into a FIB/SEM instrument equipped with an integrated fluorescence light microscope (int-FLM) for lamella preparation. While lamella milling of plunge frozen samples can be done by ablating material with the FIB from a shallow angle, milling waffle grids requires additional steps. First, the target area must be located laterally, i.e. parallel to the sample surface. Then holes, referred to as trenches, are milled around the target area with the FIB axis perpendicular to the sample surface (trench milling orientation, Figure 1 – figure supplement 3). Afterwards, the axial position of the target area is determined followed by thinning the sample from a shallow angle to obtain a lamella (lamella milling orientation, Figure 1 – figure supplement 3).

Correlating the FLM images with FIB images is difficult due to the near-featureless surface of waffle grids. For lateral targeting, a marker hole was milled in trench milling orientation (Figure 2c) into a grid square containing a target area identified previously in the FLM. Marker and target area were then localized in sample top views recorded with the int-FLM (Figure 1 – figure supplement 3d) and the measured distance between them was used to guide trench milling (Figure 1 – figure supplement 3e).

To ensure that the cell junction is contained within the final lamella with a thickness of only a few hundred nanometers, it must also be targeted in the axial dimension, i.e. within the thickness of the sample. For plunge frozen samples, 3D-correlative light microscopy and FIB milling allows axial targeting by using cryo-FLM z-stacks^12^. This approach, however, is difficult to apply to thick samples, because of poor axial resolution, depth dependent aberrations and difficulties in registration between 3D-stacks and FIB images of HPF samples. We applied two different imaging modalities for axial targeting. First, cellular components like cell walls, membranes or organelles can be visualized by SEM block-face imaging^34^, if they are exposed at the FIB milled surface after trench milling (Figure 2d, Figure 1 – figure supplement 3c). Second, when the target area is buried beyond the surface, the int-FLM can be used in the “FIB-view” mode (Capitanio *et al*., in preparation). Here, int-FLM images are not acquired as top views but with the stage in lamella milling orientation (Figure 1 – figure supplement 3b) from the same shallow angle at which the lamella is thinned to its final thickness (Figure 2e).

Next, the lamella was further thinned in several successive steps (Figure 2 – figure supplement 2). At a thickness of 6 µm a notch similar to reference^21^ but larger was milled to allow for movements of more than 1 µm to prevent lamella bending in the type of samples studied here (Figure 2 – figure supplement 2b). Afterwards, the lamella was thinned to its final thickness of 100-300 nm (Figure 2 – figure supplement 2c) and imaged in a TEM. Lamella overviews reveal the subcellular organization in *P. patens* protonemata unperturbed by crystalline ice (Figure 1b, Figure 3d).

**Figure 3.**
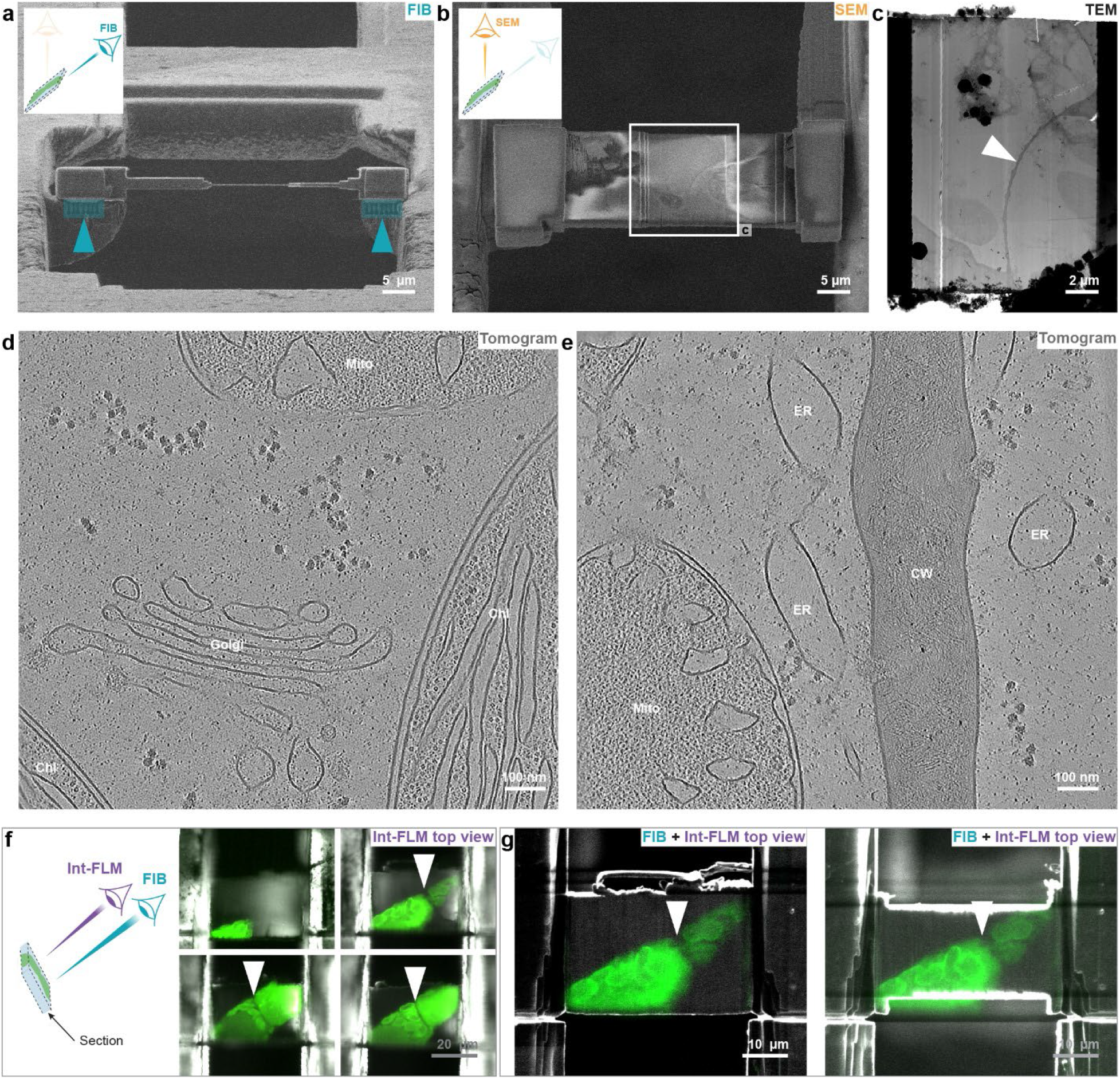
Serial Lift-Out with double sided attachment from below and section trimming for *P. patens* protonemata. **a-c**, Double-sided attached lamellae with the attachment from below. FIB and SEM images are shown in **(a)** and **(b)**, respectively. The attachment sites are marked in blue. The white rectangle in **(b)** marks the field of view in **(c)**. **c**, TEM overview image of the lamella. d,e, Single slices of denoised tomograms. Chl indicates Chloroplasts, Mito Mitochondria, ER the cortical Endoplasmatic Reticulum and CW the cell wall. **f,** Four int-FLM top views of successive Serial Lift-Out sections (reflected light in gray and cellular autofluorescence in green). The junction between cells (white arrowhead), is only contained in three of them. **g**, Overlay of FIB (gray) and int-FLM top view images (green) of the top right section in **(f)** before and after section trimming in the left and right panel, respectively. Video 1: Lift-out in lamella milling orientation on a waffle grid. This approach resulted in longitudinal sections of protonemata as shown in left panel of Figure 3 – figure supplement 1,e. Video 2: Lift-out in trench milling orientation on a waffle grid. This approach can resulted in cross-sections of protonemata as shown in right panel of Figure 3 – figure supplement 1d,f. Video 3: Lift-in with single-sided attachment. At this point we had serious recontamination problems in our FIB-SEM instrument. In this case, the marker line was simply milled into the contamination that accumulated on the pin over time. An example for a single-sided attached lamella is shown in Figure 5a,b. Video 4: Lift-in with double-sided-attachment from the side. Video 5: Lift-in with double-sided-attachment from below. An example for a lamella with double-sided attachment from below is shown in Figure 3a-c. Video 6: *Physcomitrium patens* protonema cell. Example of a tilt-series with the corresponding, denoised tomogram. A single slice of the tomogram is shown in Figure 3d. Video 7: Lift-out in lamella milling orientation on an in-carrier frozen sample. Video 8: Lift-out in trench milling orientation on an in-carrier frozen sample.

### Optimization of Serial Lift-Out for lamella quality

The cell junctions we targeted in this work have a diameter of 15-25 µm. The process of waffle milling results in a single lamella of a few hundred nanometers in thickness. Thus, about 98% of the targeted volume is lost. Additionally, as targeting and trench milling are time-consuming processes, the yield of a single lamella per site renders the technique low throughput. An alternative to waffle milling is Serial Lift-Out^22^. Here, the entire target area is extracted by lift-out (Video 1 and 2) and cut into several sections that are attached to a receiver grid. Those sections are then fine milled, resulting in multiple lamellae sampling the target area, increasing the throughput and the contextual information obtained^22^.

We explored both attachment methods proposed in the original paper: single-sided and double-sided attachment. For the single sided attachment, redeposition milling was used to attach only one side of a section to a pin (Figure 3 – figure supplement 2, Video 3). In our hands, this was the simplest attachment method as it required less precise maneuvering of the lift-out system, but the resulting lamellae often bent during fine milling (Figure 3 – figure supplement 2f), especially when thinned below 200 nm as described previously^22^. Lamella loss during the transfer between microscopes was also more frequent (Supplementary Figure 6). Furthermore, lamellae were unstable during data acquisition, often resulting in poor data quality.

For the double-sided attachment, the width of the lift-out block was adjusted to fit precisely between two grid bars of a 400×100 mesh grid. Attachment of the section on both sides (Figure 3 – figure supplement 3, Video 4) prevents bending during fine milling and results in lamellae which are stable during microscope transfers and data acquisition. This approach, however, requires more precision with the micromanipulator system and user experience.

In the context of the present study, we adapted the double-sided attachment method to focus on lamella quality rather than quantity. To this end, we milled plateaus into grid bars onto which the lift-out block could then be placed. The sections were then attached to the plateaus from below (Figure 3a, Figure 3 – figure supplement 4, Video 5). While milling the plateaus increases the expenditure of time (15-25 min per section), it leads to robust attachment and is easier to learn for lift-out beginners because it requires less precision concerning the dimensions of the lift-out block. The reduced precision requirement also allows Serial Lift-Out to be performed on FIB systems with stage instabilities or with imperfections in the FIB optics. For all three attachment methods, we obtained vitreous lamellae (Figure 1b, Figure 3c) and recorded tomograms revealing the subcellular organization of *P. patens* with an unprecedented level of detail (Figure 3, Video 6). The seemingly discontinuous membranes in Figure 3d,e are caused by the missing wedge, a known imaging artifact in cryo-ET ^35^.

Generating homogeneously thin lamellae is challenging, especially if they are larger than 10-20 µm along the milling direction. To improve lamella quality, we included an additional step to the Serial Lift-Out workflow, namely section trimming. After the attachment, int-FLM top views were acquired for all sections to check which of them contained the target area (Figure 3f). This also allowed to identify regions of the sections with no features of interest which were removed by FIB milling in trench milling orientation (Figure 3g), minimizing the depth of the lamella in the thinning direction. During the lift-out process, the front of the lift-out block becomes damaged and uneven due to repeated ion exposures. At least 2 µm of the section front were also removed to create a new, smooth front. This trimming step makes fine milling faster because less material must be ablated and allows to generate homogeneously thin lamellae as the smooth front prevents curtaining^36^.

### Lift-out in different stage orientations

On waffle grids of *P. patens*, the axis of the cylindrical protonemata cells is always roughly parallel to the grid surface. Thus, on-grid waffle milling and lift-out with the stage in the lamella milling orientation (Figure 3 – figure supplement 1 left panel, Video 1) will inevitably yield sections roughly parallel to this axis (longitudinal sections). Performing lift-out in the trench milling orientation on cells with the protonemal axis oriented roughly parallel to the lift-out direction, cross-sections can be obtained (Figure 3 – figure supplement 1 right panel, Video 2), analogous to *Caenorhabditis elegans* L1 larval cross-sections in reference^22^. Thus, it is possible to address samples which can only be frozen in distinct orientations from different directions, which is crucial to study subcellular structures with defined orientation within cells.

### In-carrier high-pressure freezing

During waffle freezing, *P. patens* protonemata can be damaged for two reasons. First, the filaments are larger than the squares of the EM-grids used and cells running across grid bars will be squeezed. Second, often multiple cell layers were pressed into the 25 µm high well formed by the grid bars which is not high enough to accommodate them without compression. To avoid this mechanical damage, we high-pressure froze protonemata directly in HPF carriers with a 2 mm wide and 100 µm deep cavity.

While the procedure of in-carrier high-pressure freezing is fast, it comes with additional challenges. First, samples cannot be screened easily in a FLM because carriers do not fit into the commercial shuttle. Thus, montage overviews were recorded with the int-FLM (Figure 3 – figure supplement 5a). In our systems, this takes longer and yields poorer quality images due to widefield imaging and the low numerical aperture of the objective. Nevertheless, the obtained montage overviews were sufficient to identify target areas. Trench milling aided by int-FLMs was then performed as described in the previous section (Figure 3 – figure supplement 5b).

Second, lamellae must be prepared by lift-out methodologies as trench milling through the carrier is impractical. Additionally, the lift-out block must be detached from the bulk also from below and not only on all four sides as for waffle grids. This requires milling an undercut after trench milling in lamella milling orientation (Figure 3 – figure supplement 5c, Video 7 and 8).

Third, in-carrier HPF samples are thicker than typical waffle grids. Thus, the axial targeting via SEM block-face imaging or FIB-view imaging with the int-FLM is crucial to ensure that the undercut is milled below the target area and that it is extracted completely (Figure 3 – figure supplement 5c). Apart from these technical complications, lift-out and sectioning was then performed as for waffle samples, with the added benefit that in-carrier HPF potentially allows to vitrify thicker samples.

### Subtomogram averaging of rubisco

In order to test if structural details are preserved throughout the presented workflows, we subjected rubisco^37^, an abundant enzyme in chloroplasts, to subtomogram averaging^38^. We acquired a small dataset composed of 10 tomograms on *P. patens* chloroplasts, picked rubisco positions by template matching and computationally aligned and classified subtomograms. Figure 4a shows an example of a chloroplast tomogram (Video 9), and Figure 4b the corresponding 3D rendered image of segmented chloroplast membranes with an intermediate, low-resolution rubisco average pasted at the determined particle positions (see supplementary material). We docked a model of an octamer of large rubisco subunits predicted by AlphaFold 3^39^ into the final average of about 6000 subtomograms (Figure 4c). The α-helices of the predicted model fit well into the experimental density (Figure 4d) indicating that we attained an average with subnanometer resolution (Supplementary Figure 4, FSC_@0.143_ = 0.79 nm, FSC_@0.5_ = 0.95 nm).

**Figure 4.**
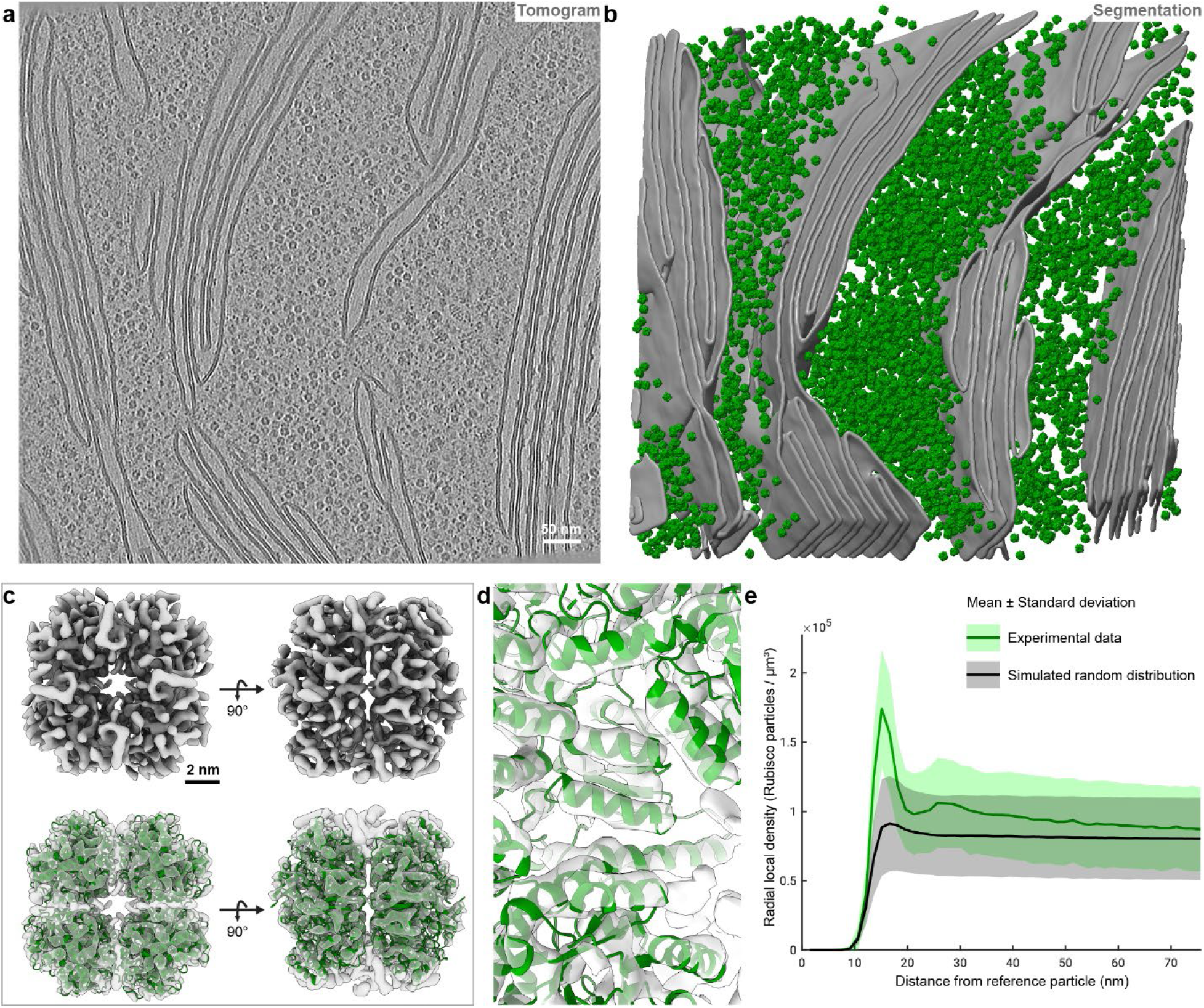
Subtomogram averaging of rubisco in chloroplasts of *P. patens* protonemata. **a**, Slice of a denoised tomogram recorded on a chloroplast (Video 9). **b,** Three-dimensional rendered representation with segmented chloroplast membranes (gray) and the rubisco subtomogram average (green) pasted at the positions determined after template matching and subtomogram classification. **c,** Rubisco subtomogram average in isosurface representation (top row) and an octameric model of large rubisco subunits predicted by AlphaFold 3 docked into the average (bottom row). **d,** Close-up view of the average shown in (c) revealing that predicted α-helices fit well into the experimental density map. **e**, The local density of rubisco particles as a function of the distance from the reference particle. Green line and area are the average of experimental density ± one standard deviation, respectively, obtained from 10 tomograms. Black line and gray area are the mean ± one standard deviation obtained for a simulated random particle distribution. Video 9: *Physcomitrium patens* protonema chloroplast. Example of a tilt-series with the corresponding, denoised tomogram. A single slice of the tomogram is shown in Figure 4a.

Based on the determined positions of rubisco complexes within tomograms, we calculated the local rubisco density which has a peak at 16 nm in contrast to a simulated random distribution (Figure 4e). This indicates that rubisco is not randomly distributed within *P. patens* chloroplasts similar to previous findings in the pyrenoid of the green alga *Chlamydomonas reinhardtii*^40^. These results demonstrate that subnanometer structural details are preserved throughout the presented workflows and that it is now possible to perform *in situ* structural biology studies in multicellular plants.

### The high-pressure freezing workflow applied to other tissues and plants

To demonstrate the application of the HPF methods presented to other tissues and plant species, we further used it on phyllids, the leaf-like structures formed by *P. patens*. Small pieces of phyllids were dissected and frozen with the Waffle Method. The samples were screened to avoid investigating areas which have been damaged during dissection or freezing (Figure 5 – figure supplement 1a-c). We were able to obtain vitreous lamellae and tomographic data from this tissue (Figure 5a,b, Video 10).

**Figure 5.**
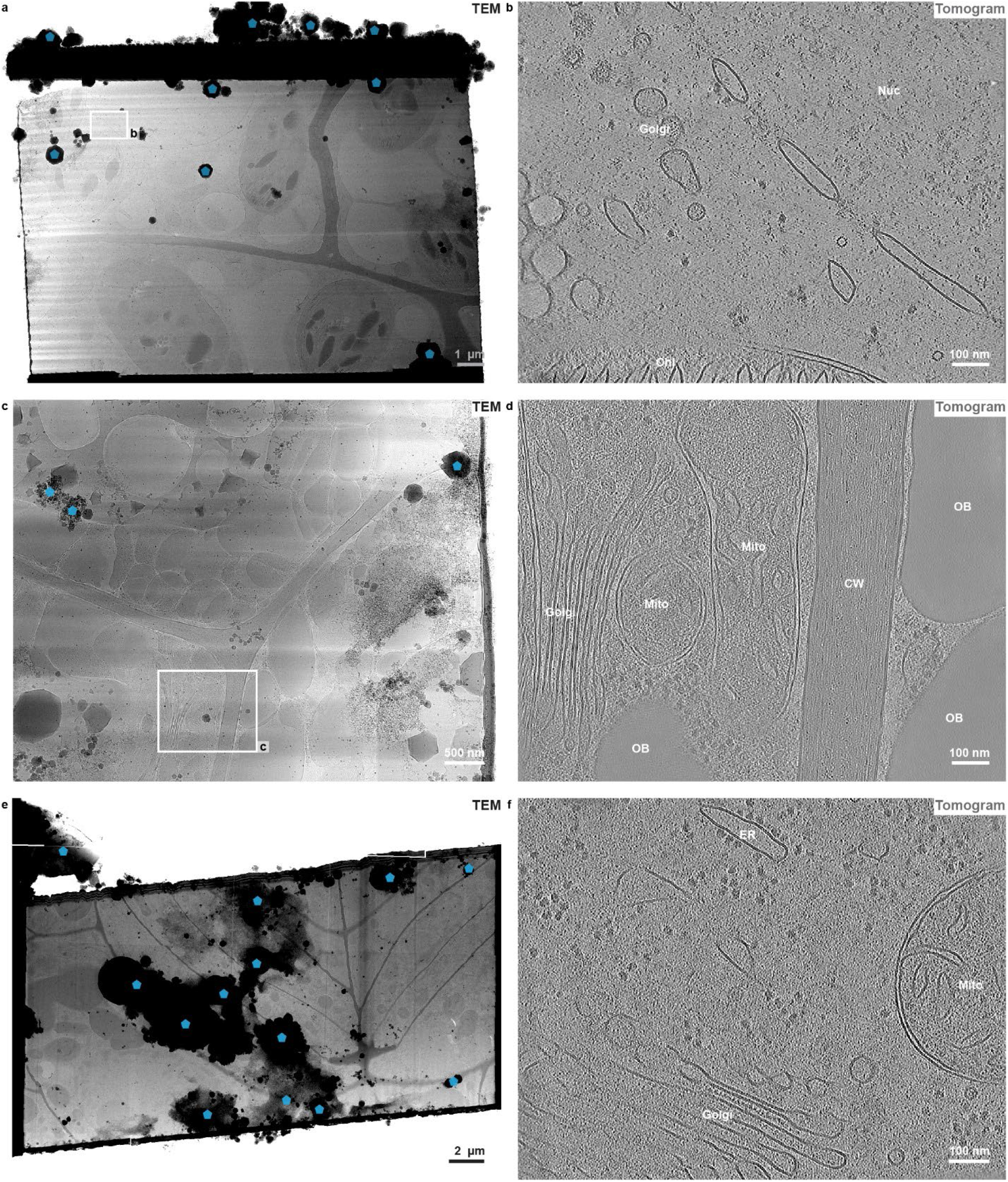
The cryo-ET workflow applied to other tissues and species. **a,b**, Physcomitrium patens phyllid sample. **a,** TEM overview image of a single-sided attached lamella. **b,** a single slice of a tomogram recorded on the nuclear envelope (Video 10). **c,d,** Arabidopsis thaliana embryo sample. **c,** Low magnification TEM image of a lamella. **d,** Single tomographic slice of the cell wall between cells (Video 11). **e,f** Salt gland of Limonium bicolor. **e,** TEM overview image of a double-sided attached lamella. **f**, Single slice of a tomogram (Video 12). Chl indicates Chloroplasts, Mito Mitochondria, OB oil bodies, Nuc the nucleus, CW the cell wall and ER the Endoplasmatic reticulum. All tomographic slices are from computationally denoised tomograms. In (a), (c) and (e) some ice crystals contaminating the lamella surface which were introduced during transfers between microscopes are marked by blue transparent pentagons. Video 10: *Physcomitrium patens* phyllid. Example of a tilt-series with the corresponding, denoised tomogram. A single slice of the tomogram is shown in Figure 5b. Video 11: *Arabidopsis thaliana* embryo. Example of a tilt-series with the corresponding, denoised tomogram. A single slice of the tomogram is shown in Figure 5d. Video 12: *Limonium bicolor* salt gland. Example of a tilt-series with the corresponding, denoised tomogram. A single slice of the tomogram is shown in Figure 5f.

Next, we adapted the waffle workflow for *Arabidopsis thaliana* embryos. While the embryos easily fit into grid squares, they are with about 50 µm thicker than the 25 µm high bars of EM-grids used here (Figure 5 – figure supplement 1d,e). To avoid mechanical damage, an additional spacer ring was added onto the grid before waffle freezing, which increased the thickness of the waffle sample to around 75 µm and prevented squeezing of the embryos in the process (Figure 5 – figure supplement 1d,e). *A. thaliana* embryos were successfully vitrified, and tomographic data acquired (Figure 5c-d, Video 11).

As a last study subject, we high-pressure froze the first true leaves of *Limonium bicolor* (Figure 5 – figure supplement 1f) in the 300 µm deep cavity of HPF carriers. We targeted the blue autofluorescent salt glands (Figure 5 – figure supplement 1g-h) again aided by the int-FLM and performed Serial Lift-Out. A TEM overview of a salt gland section and a tomographic slice are depicted in Figure 5e and f, respectively, revealing its cellular architecture in a near-native state (Video 12). These results illustrate that the presented HPF methods are broadly applicable.

## Discussion

Compared to the “standard” cryo-ET workflow, comprising plunge-freezing, on-grid lamella milling and data acquisition, working with high-pressure frozen samples is technologically more demanding and has a lower throughput and success rate. Hence, if a sample can be vitrified by plunge-freezing, it still is the method of first choice. We must assume, however, that most plant tissues are too thick to be vitrified by plunge-freezing and must instead be high-pressure frozen.

Depending on the size of the sample, there are several high-pressure freezing variants. For the Waffle Method without spacer ring, the thickness of the EM-grid defines the maximum thickness of the sample. While standard EM-grids have a thickness of 20-25 µm, thicknesses between 8-100 µm are commercially available. Additionally, the shape (e.g. square, hexagon, hole and slot) and the size of the mesh can be chosen to match the shape of the tissue. For samples exceeding 100 µm thickness, spacer rings can be added, as exemplified here on *A. thaliana* embryos. The advantage of waffle grids are that they allow on-grid lamella preparation and no undercut is required when performing lift-out. Samples with diameters of up to 2 mm and a maximum thickness of 200-300 µm can potentially be high-pressure frozen in HPF carriers without risking squeezing damage by grid bars. In this case, on-carrier lamella preparation is currently technologically impractical and lift-out methodologies are required.

Vitrification is fundamental for preserving the sample in a near-native state and has a strong impact on the cryo-ET data quality and subsequently subtomogram averaging. We obtained data from clearly vitrified, thus apparently structurally intact samples in many cases. The success rate, however, was very sample-dependent. All our *A. thaliana* embryo and *L. bicolor* samples were vitreous (N = 4 and N = 1, respectively), perhaps due to the cells, which are densely packed with cellular components and the absence of large vacuoles. For *P. patens*, the success rate was 47% (N = 45, Supplementary Figure 6a). Occasionally, even cells from within the same grid square differed in their vitrification state. We applied similar methods to the trichomes of *Nicotiana benthamiana* and observed large ice crystals in all of them (N = 10). This illustrates that obtaining consistent vitrification results with high-pressure freezing is still a major challenge. Moreover, certain samples require harvesting or dissection before being manually mounted in HPF carriers, which inevitably damages or stresses the cells and may result in artifacts. Hence, the preparation of samples for HPF should be as quick as possible while minimizing the perturbation of the system under investigation. It would therefore be beneficial to develop new protocols or even methods, that are largely user independent or completely automated, highly reproducible and capable of even vitrifying larger and thicker tissues^41,42^. Currently, most fully intact tissues remain only accessible by conventional electron microscopy employing chemical fixation, if the dissection to fit the sample into the small volume of HPF carriers is omitted.

Additionally, HPF samples demand a targeted approach, since the cells are not exposed on the sample surface but submerged in a layer of vitreous buffer. Usually, the localization of target areas requires fluorescent markers or autofluorescence. For lateral targeting in the x-y-plane of the sample, grids were screened in a stand-alone cryo-FLM to identify suitable grid squares. Next, int-FLMs proved practical to guide trench milling. To ensure that the target area is contained within the final lamella or lift-out block, its axial position (location within sample thickness) must be determined. SEM block-face imaging after trench milling is one option. It is only applicable, however, if sufficiently contrast-rich features, like membranes, in the vicinity of the target area are exposed at the block-face. The second option is FIB-view imaging which can be applied if a fluorescently labelled or autofluorescent target is discernible in int-FLM images.

On-grid waffle milling has the lowest methodological entry barrier and does not require lift-out. It is faster for screening several locations per grid and assessing the sample quality than Serial Lift-Out. At the same time, it exhibits a lower throughput by obtaining only a single lamella per targeted area. Additionally, fine milling for our waffle samples must be done at relatively high stage tilts (20°) to fully access the target area and it is often difficult to remove all material below lamellae. Both factors restrict the angular range for tomographic data acquisition and, in turn, compromises data quality. For large target areas, like in our case the junction between cells, Serial Lift-Out is the preferred method. While it requires some training and a lift-out system, after sectioning, fine milling is faster and more straightforward than for waffle milling and can be performed at shallow angles. The full ±60° tilt range for cryo-ET, can therefore be accessed.

For plant samples, single-sided attachment is the fastest lift-out approach requiring least user precision. But smaller lamella dimensions, potential bending and instabilities during data collection must be considered. In comparison, double-sided attachment yields lamellae that are easier to thin homogeneously without bending and produce higher quality data. For the double-sided attachment, we tested two methods. The side-attachment (i), as described originally in reference^22^ needs little modifications of the receiver grid and consequently requires less expenditure of time. Yet, fitting the lift-out block precisely between the grid bars allows only for an error of a few hundred nanometers. Hence, it requires most user expertise, a mechanically very stable stage and lift-out system as well as well-aligned ion optics. The attachment from below (ii) is more time-consuming due to the additional generation of plateaus but adjusting the size of the lift-out block and placing it on the plateaus tolerates manipulation errors of a few micrometers. Therefore, the attachment from below is likely easier to adopt for lift-out novices.

The additional trimming step introduced here focuses on obtaining highest lamella quality over quantity. First, int-FLM top views of Serial Lift-Out sections confirm that the target area is captured within followed by removal of regions that do not contain material of interest in trench milling orientation. This minimizes the lamella size in the fine milling direction and centers the target between smooth fronts which is important to obtain homogeneously thin lamellae during the subsequent fine milling in lamella milling orientation.

To further increase the throughput and reduce the experience defined entry barrier, automation is important. Since we explored multiple samples which all behaved differently, all milling steps were performed manually. The whole lamella preparation workflow, however, is per-se very repetitive and time-consuming. Several software tools already exist^18,43–45^ which can be used for on-grid waffle milling and fine milling of lift-out sections. Yet, robust automated protocols for targeted trench milling, lift-out and sectioning must still be developed.

In summary, we applied various HPF and lamella preparation methods to different plant species and tissues. We discuss their advantages and disadvantages and provide detailed descriptions of the procedures that worked best for plant tissues. We demonstrate that subnanometer structural details are preserved in the process and that meaningful spatial information can be retrieved using rubisco as a model case. Furthermore, the described workflows were successfully applied to study Plasmodesmata in *P. patens* ^46^. Hence, the presented toolbox allows for the first time to do *in situ* structural biology in plant tissues and can likely be adapted to other multicellular systems.

## Acknowledgement

We thank Dustin Morado, Tillman Schäfer, Daniel Bollschweiler, Dominik Hrebik and Hui Guo for support with the TEMs, the lab of Ben Engel for many fruitful discussions, Iuliia Iermak for help with 3D rendering and Florian Beck for invaluable input on data processing. The *Limonium bicolor* Kuntze (Plumbaginaceae)^47,48^ seeds were kindly provided by Fang Yuan and Bao-Shan Wang from Shandong Normal University. This work received funding from the European Research Council (ERC Grant agreements ‘SymPore’ no. 951292). Open access funding was provided by Max Planck Society.

## Author Contributions

Conceptualization: WB, JMP, WBF, RS, WXS

Methodology: OHS, COJK, SK, AB, CC, JB, MD, MP, JM, ML

Software: MP

Investigation: MP, MD, PX, ML, JM, COJK, OHS

Resources: WB, JMP, WBF, RS, WXS, MM, NSKJ,

Writing—original draft: MP, MD, OHS, COJK, PX

Writing—review & editing: All authors

Visualization: MP, MR, MD, PX

Supervision: WB, JMP, WBF, RS, WXS

Project administration: WB, JMP, WBF, RS, WXS, MP, MD, PX

Funding acquisition: WB, JMP, WBF, RS, WXS

## Data availability

The rubisco subtomogram average with the corresponding dataset (movie frames and meta data) have been deposited to EMDB under the accession code EMD-53132 and EMPIAR-12644, respectively. Examples of tomograms recorded on *P. patens* phyllids, *A. thaliana* embryo and *L. bicolor* have been deposited to EMDB under accession codes EMD-53142, EMD-53134 and EMD-53136, respectively.

## Methods

Plunge-frozen and high-pressure frozen samples were prepared with a Vitrobot Mark 4 (Thermo Fisher Scientific, Waltham, Massachusetts, USA) and a Leica EM-ICE (Leica Microsystems, Wetzlar, Germany), respectively. Three gallium cryo-focused ion beam (FIB) devices were used: (i) a Scios (Thermo Fisher Scientific, Waltham, MA, U.S.A) equipped with an integrated fluorescence microscope (int-FLM: Meteor (Delmic, Delft, Netherlands), MPLFLN 50× objective (Olympos), NA = 0.8, working distcance = 1 mm), (ii) an Aquilos 1 (Thermo Fisher Scientific) with an int-FLM (Meteor (Delmic), LMPLFLN 50x objective (Olympos), NA = 0.5, working distance = 10.6 mm) and an Aquilos 2 (Thermo Fisher Scientific). All FIBs were additionally equipped with a needle micromanipulator (EasyLift, Thermo Fisher Scientific) for lift-out experiments. The procedures described in the following (including stage orientation and milling pattern orientation) are only correct for the standard FIB/SEM geometry employing a shuttle with a pre-tilt of 45° (Figure 1 – figure supplement 3a) and scan rotation for FIB and SEM of 180°.

**Figure 1 – figure supplement 1:**
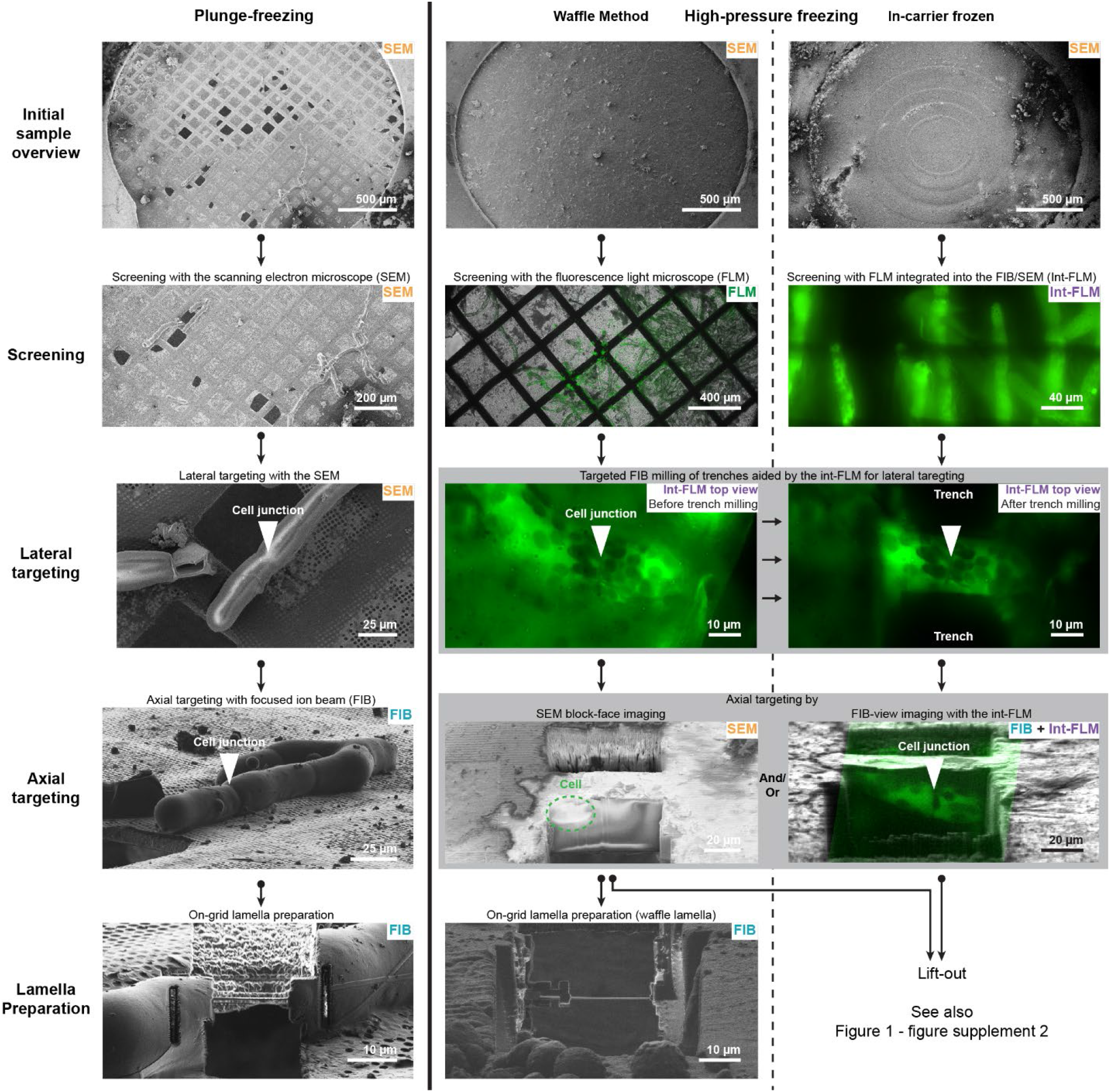
Overview of the critical workflow steps for *P. patens* protenemata. The left panel shows the workflow for plunge frozen tissue which was not vitrified. Milling lamellae at cell-junctions does not require targeting aided by cryo-fluorescence light microscopy (cryo-FLM). If a specific structure inside the cell is targeted, however, cryo-FLM can be crucial for working with plunge frozen samples as well. The middle and right panel illustrate the high-pressure freezing (HPF) workflows. The Waffle Method is depicted in the middle. After targeting, there are two options for lamella preparation. On-grid lamella milling or Lift-Out. The right panel shows the in-carrier HPF workflow. Here, lift-out is the only way to obtain lamellae for cryo-ET. The employed lift-out procedures are summarized in Figure 1-figure supplement 2. The shown imaging modalities with respect to the FIB/SEM stage and sample orientations are depicted in Figure 1 – figure supplement 3. Note that the excitation wavelengths and emission filters used for FLM imaging are different from those used for int-FLM imaging. Therefore, the fluorescence signals appear different. Ideally, for most reliable targeting, excitation and emission should be the same for all FLM imaging modalities.

**Figure 1 – figure supplement 2:**
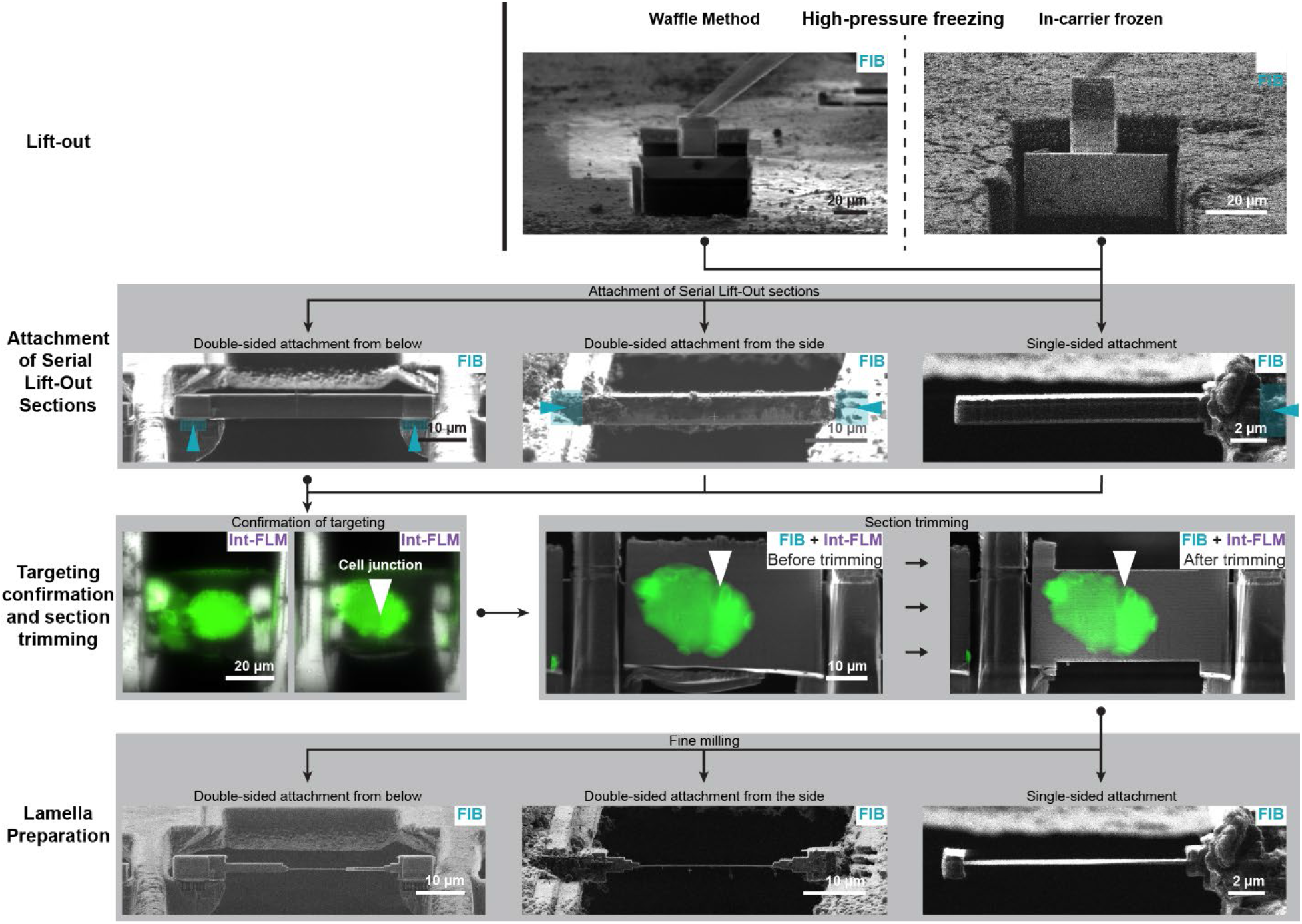
Overview of the used lift-out procedures. After the lift-out from waffle grids or in-carrier frozen samples, the lift-out block is sectioned. The section can be attached either on a single side, from two sides or on two sides from below. Attachment sites are indicated by blue rectangles and arrowheads. Afterwards, it was confirmed that target areas are contained in the sections, followed by section trimming and fine milling of sections to prepare lamellae with a thickness of a few hundred nanometers.

**Figure 1 – figure supplement 3.**
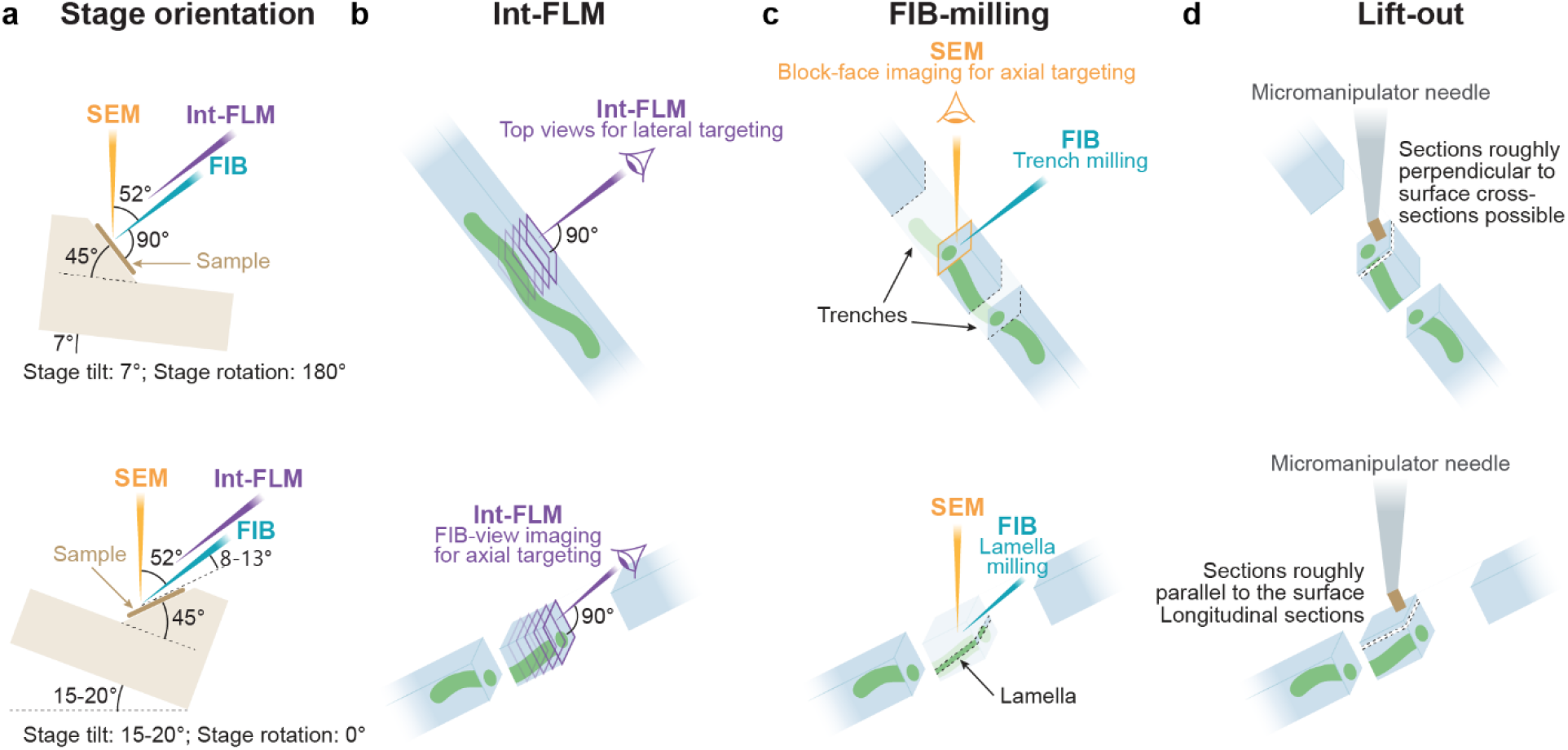
The different stage orientations used during lamella preparation. Top and bottom row show the trench milling and lamella milling orientation, respectively, for the geometry in FIB/SEM instruments produced by Thermo Fisher Scientific using a shuttle with 45° pre-tilt. **a**, The shuttle with respect to stage tilt, stage rotation and the relative location of the FIB and SEM beams. **b**, Use of the integrated fluorescence light microscope (int-FLM). In trench milling orientation (top panel), the int-FLM records sample top views for lateral targeting and in lamella milling orientation (bottom panel) it can be used in FIB-view mode for axial targeting. **c**, FIB milling. In trench milling orientation (top panel), the ion beam is normal to the sample surface and used to mill trenches above and below the target area. In Lamella milling orientation (bottom panel), the FIB is used to thin the lamella to a thickness of about 100-300 nm from a shallow angle. **d**, Lift-out. In trench milling orientation (top panel), the lift-out results in sectioning planes roughly normal to the sample surface and cross-sections of protonemata can be obtained. In lamella milling orientation (bottom panel), sectioning planes are roughly parallel to the sample surface resulting in longitudinal sections of protonemata.

### *Physcomitrium patens* culture/growth

*Physcomitrium patens* (Hedw.) Gransden ecotype was cultivated on solid BCDAT medium overlayed with cellophane sheets at 25°C and 18h white light/6h dark cycle as described in^49^. Protonemata were harvested after 2 weeks. For plunge-freezing, the tissue was blended in liquid BCDAT medium, allowed to regenerate for 2 – 4 h and then used for cryo-sample preparations. Gametophores were harvested after 4 weeks. For cryo-fixation, individual phyllids or the gametophore tip containing multiple phyllids were cut off, transferred to liquid BCDAT medium and immediately applied to EM-grids or HPF carriers.

### Plunge-freezing and FIB milling of *P. patens* tissues

BCDAT suspended tissues were applied to glow-discharged EM-grids (Quantifoil Cu 200 mesh, holy carbon film R2/1) and transferred to a Vitrobot Mark IV (Thermo Fisher Scientific), temperature set to 25°C and humidity control switched off. Blotting was performed with a Teflon sheet from support film side and a filter paper from the reverse side for 10 s with a blot force 10, before plunging into a liquid ethane-propane mixture^50^.

Additionally, different cryoprotectants were tested. Either by culturing *P. patens* on solid BCDAT medium containing cryoprotectants, by incubating *P. patens* after blending in liquid BCDAT medium containing cryoprotectants, by adding 3 µL BCDAT containing cryoprotectants to the EM-grid right before plunging or by combinations of these methods. The different methods with the tested concentrations are listed in Table 1. The use of cryoprotectants either did not prevent the formation of crystalline ice or induced plasmolysis (data not shown) and sometimes even dehydrated the tissue (Figure 1 – figure supplement 4f).

Grids were clipped into cartridges with cutouts (Thermo Fisher Scientific) and loaded into the FIB/SEM instrument. A layer of metal organic platinum was applied for 40 s using the gas injection system (GIS) with the stage in the deposition position (here, stage tilt = 25°, stage rotation = 0°, stage z = 12.5 mm, the GIS deposition position may be different for other instruments). Lamellae were milled with the stage in lamella milling orientation (stage tilt = 15°, stage rotation = 0°, Figure 1 – figure supplement 3c). This was done in several steps reducing the thickness to 6 µm, 3 µm and 1.5 µm at currents of 1 nA, 0.5 nA and 0.3 nA, respectively. Fine milling was performed at 50 pA, aiming for a final lamella thickness of 200-250 nm. Finally, the lamella was polished at 30 pA at a stage tilt of 14° and 16° only from the bottom and the top, respectively (Figure 1 – figure supplement 4a-d).

**Figure 1 – figure supplement 4.**
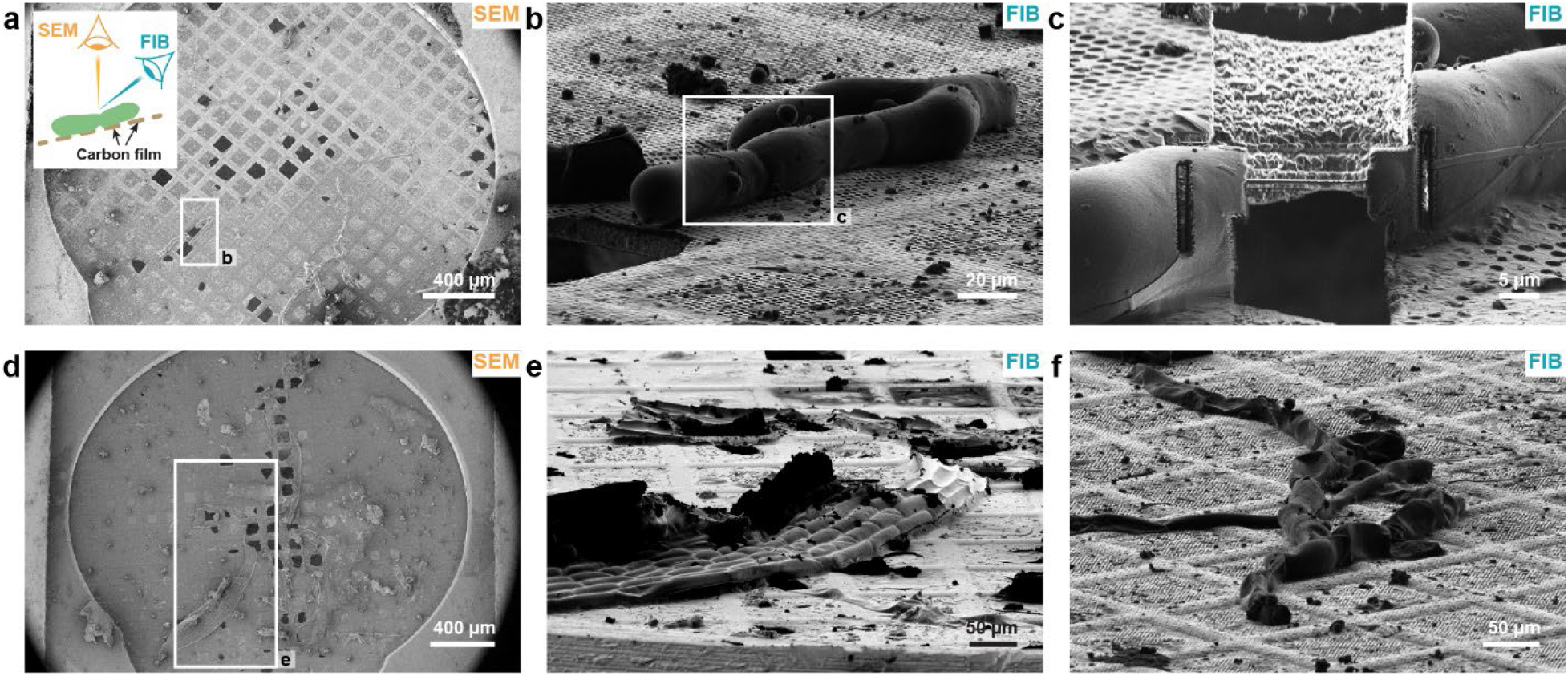
Plunge-frozen *P. Patens* tissues and the use of cryoprotectants. **a-c**, Plunge frozen protonemata. **a**, SEM overview image of a grid. **b**,**c**, FIB image before and after lamella milling, respectively. **d**,**e**, SEM and FIB images of plunge frozen phyllids, respectively. **f**, Protonemata grown on medium and plunged in medium containing 100 mM proline as cryoprotectant. The cells are dried out.

### The waffle workflow for *Physcomitrium patens* protonemata

Grids were high-pressure frozen and milled using an adapted workflow of the Waffle Method^21^ similar to the procedure described in^22^. The FIB milling and imaging step described in the following were performed in two main stage orientations. The lamella milling orientation (stage tilt = 15-20°, stage rotation = 0) and trench milling orientation (stage tilt = 7°, stage rotation = 180°). A scheme of the stage orientations is depicted in Figure 1 – figure supplement 3. More details about the workflow can be found in the step-by-step protocol (Pöge *et al.*, protocol was accepted for publication).

### High-pressure freezing of waffle grids

As described in^22^, before the experiment, 6mm Type B carriers (300 µm cavity on one side and flat on the other, Leica Microsystems, Wetzlar, Germany) were incubated overnight in a 1:1 mixture of 4 M NaOH and commercial bleach, rinsed 3x with water, 1x with Acetone and dried. Next, the carriers were dipped into a cetyl palmitate solution (0.5-1% (m/v) in diethyl ether), excess of liquid was removed with a filter paper and the carriers were dried on a filter paper with the flat side facing up. Commercially available Copper 50, 75 or 100 mesh grids with continuous support film (Carbon coated Formvar, Graticules Optics Ltd, Tonbridge, UK) were used to support the sample. As freezing buffer, a 1:1 mixture of growth medium and 40% (m/v) Ficoll 400 (Roth, Karlsruhe, Germany) was prepared. A 2 µL droplet was applied to the flat side of a carrier, a grid with the support film facing down placed on it and the excess of buffer removed with a small piece of Whatman No 1 filter paper (Whatman, Maidstone, UK) until the grid was lying flat on the carrier with a thin layer of buffer between them. 3 µL buffer were added onto the grid and distributed with the pipet tip until the whole grid surface was covered. Air bubbles were removed with tweezers. Next, a small patch of *P. patens* protonemata was picked from a plate and placed in the center of the grid. A second carrier was placed on top, flat side down, and the whole sandwich tightly squeezed together for 10 s followed by immediate high-pressure freezing. The resulting waffle grids were separated from the carriers, clipped into cartridges with cutouts (Thermo Fisher Scientific) and stored until use in liquid nitrogen (LN_2_).

### Cryo-fluorescence light microscopy

Waffle grids have a near featureless surface in SEM and FIB images (Figure 2 – figure supplement 1a). Fluorescence microscopy is therefore vital for targeted lamella preparation. For initial screening, clipped grids were loaded into a confocal laser scanning microscope equipped with cryo-stage (Leica TFS SP8, Leica Microsystems, 50x Air objective: Leica Objective No. 506520, NA 0.9, two hybrid detectors and one photomultiplier tube). We always detected the reflected light (excitation wavelength: 488 nm; emission filter: 480-496 nm), the cellular autofluorescence of the tissue (excitation wavelength: 488 nm; emission filter: 500-750 nm) and the transmitted light. Dependent on the sample thickness, we adjusted the laser power to have enough signal while not overexposing the detectors. Initially, a grid map was acquired (Figure 2 – figure supplement 1b) with a spiral scan (512×512 pixel per tile image, pin hole size = 600 µm, pixel size = 580 nm). Promising grid squares containing cell junctions were subsequently imaged by acquiring z-stacks encompassing the whole sample thickness (2048 x 2048 pixel per image, pin hole size = 125 µm, pixel size = 140 nm, z step size = 1 µm, 25-50 steps). Images were recorded using the LAS X software (Leica Microsystems, Wetzlar, Germany). The FLM images displayed in the figures are exclusively single slices of z-stacks and no maximum intensity projections. Squares with several layers of protonemata were ignored because the ∼25 µm high wells of the used grids can only accommodate a single cell layer risking squeezing damaged during freezing (Figure 2 – figure supplement 1d). Here we used a confocal laser scanning microscope, but a standard widefield system equipped with a cryo-stage could also be used for this screening step.

### Lateral targeting and trench milling

Grids were loaded into a FIB/SEM instrument with integrated fluorescence light microscope (int-FLM). A protective layer of metal-organic platinum was applied for 45 s using the GIS with the stage in trench milling orientation (stage tilt = 7°, stage rotation = 180°, Figure 1 – figure supplement 3). With the stage in trench milling orientation the coincidence point of FIB and SEM was found on a grid square which contained a target area and a marker pattern (cross-section, xyz: 8 x 8 x 0.5 µm) was milled at a safe distance from the target area. For lateral targeting, sample top views (Figure 1 – figure supplement 3b top panel) were recorded with the int-FLM using a reflected light (excitation wavelength = 470 nm ± 10 nm, no emission filter) to focus on the surface and to find the marker and a cellular autofluorescence (excitation wavelength = 470 nm ± 10 nm, emission wavelength = 515 nm ± 15 nm). The junctions between *P. patens* cells could be detected based on the cellular autofluorescence without additional labelling. Z-stacks were acquired (pixel size = 168 nm, 20-25 z-slices, z-step = 1 µm). The distance as well as the location of the target area with respect to the marker was determined (Figure 2 – figure supplement 1f) using the Odemis software (Delmic). Dependent on the sample thickness, the laser power was adjusted to have enough signal to detect the target area but kept at the bare minimum to not melt the sample. If both FLM and int-FLM imaging are employed for targeting, the used excitation wavelengths and emission filters should ideally be the same for both imaging modalities. This assures most reliable targeting. That is not the case here. Therefore, the fluorescence signals of FLM and int-FLM images (Figure 2, Figure 2 – figure supplement 1) appear different.

**Figure 2 – figure supplement 1.**
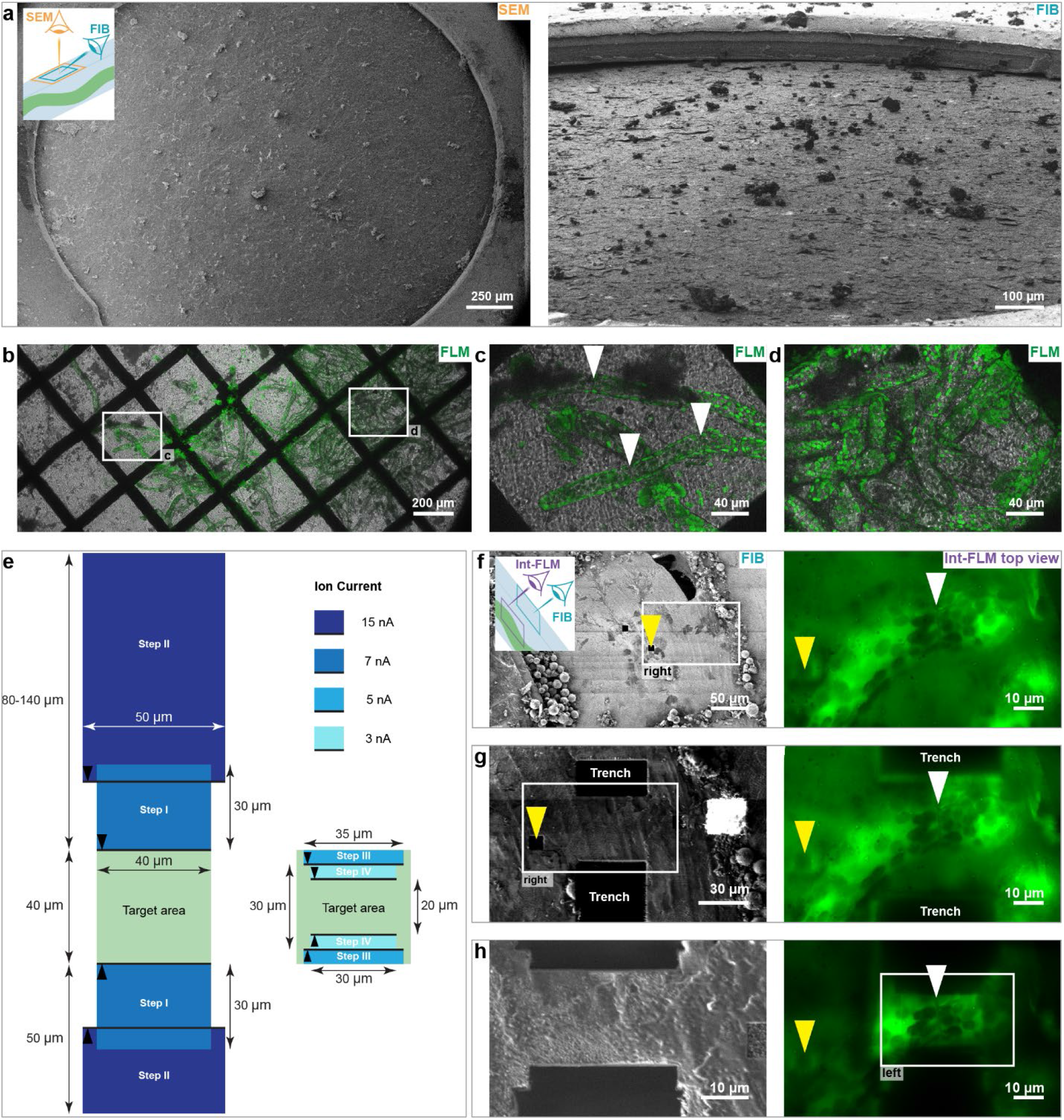
Targeted trench milling of *P. patens* protonemata aided by the int-FLM. **a**, SEM and FIB images of a waffle grid. **b**-**d**, FLM images of a waffle grid (transmitted light in gray and cellular autofluorescence in green). Grid overview in **(b).** The squares mark the field of views in **(c)** and **(d)** which are grid squares containing single and multiple cell layers, respectively. The cells in **(d)** were likely damaged during HPF. e, Scheme summarizing all trench milling steps. All trench milling steps are performed in trench milling orientation (Figure 1 – figure supplement 3). The milling direction is always towards the target area (indicated by black arrowheads) using regular cross-sections. **f-h** FIB (gray) and int-FLM top view images of cellular autofluorescence (green) for successive trench milling steps in the left and right panel, respectively. Int-FLM top views were recorded between trench milling steps to assure the target area was properly centered. First, a marker pattern was milled (yellow arrowhead) and identified in int-FLM images. From there the distance to the target area was measured (white arrowhead) in **(f).** This information was used to place the patterns for milling step I in **(e).** The results are shown in **(g)**. After trench milling the target area is centered within the block between trenches **(h). In (f)-(g)** white rectangles indicate the field of view in the second image. Note that the excitation wavelengths and emission filters used for FLM imaging in **(b)-(d)** are different from those used for int-FLM imaging in **(f)-(h).** Therefore, the fluorescence signals appear different. Ideally, for most reliable targeting, excitation and emission should be the same for all FLM imaging modalities.

Based on the measured distance of the target area from the marker, the position of trench milling patterns were defined. The drift suppression function of the instrument was activated during all trench milling steps. First, two cross-sections with an xy-size of 40×30 µm were milled above and below the target area with a distance of 40 µm at an ion current of 7 nA (step I in Figure 2 – figure supplement 1e). Next, the trenches were elongated at 15 nA to a height of 50 µm and 80-120 µm (step II in Figure 2 – figure supplement 1e), on bottom and top, respectively. The target area containing block was further thinned in two steps to an xy-size of 35×30 µm and 30×20 µm at 5 nA and 3 nA, respectively (step III and IV in Figure 2 – figure supplement 1e). Top views were acquired with FIB and int-FLM after each milling step to adjust the positioning of the patterns, assuring that the targeted area was centered after trench milling (Figure 2 – figure supplement 1g-h). Then the block-face on top was polished at 1 nA.

### Axial targeting

The images of the int-FLM allow to reliably localize the target area laterally, i.e. in parallel to the sample surface. Due to the limited numerical aperture of the objective, the wide-field epifluorescence optical setup of the int-FLM and depth dependent aberrations, the axial resolution is limited. The axial localization of the target area within the thickness of the sample is crucial to place the patterns for lamella fine milling. Two methods were used for the axial targeting: (i) SEM block-face imaging and (ii) FIB-view imaging with the int-FLM.

After trench milling, with the stage still in trench milling orientation, the SEM images the side surface of the block (Figure 1 – figure supplement 3c, top panel). Cellular structures, like membranes, exposed at the FIB milled front of the block, can be visualized with by SEM imaging, even for non-metal stained, vitrified samples^51^. Before the image was recorded, the auto-stigmator function was run on a nearby spherical contamination. The whole block-face was then centered in the SEM field-of-view, the SEM focused on the edge between grid surface and block-face and the auto brightness-contrast function run on the block-face itself. SEM images were acquired with a dwell time of 50 ns, 100 line integrations at U = 3 kV, I = 13 pA with an image size of 3072 x 2048 pixel (Figure 2d). If inherent material contrast is sufficiently strong, the approximate distance of the target area from the sample surface can then be measured in the SEM image using the FIB/SEM user interface.

For FIB-view imaging, the int-FLM was used to image the side of the block after trench milling in lamella milling orientation (Figure 1 – figure supplement 3b, bottom panel) at the angle at which the lamella will be milled. For optimal imaging, the lamella front has to be normal to the optical path of the int-FLM (Capitanio *et al*., in preparation). This allows to avoid major depth-dependent refractive-index-mismatch-induced aberrations^52^. Normal incidence is achieved by changing the stage tilt angle during trench milling steps III-IV (Figure 2 – figure supplement 1e). The required stage tilt depends on the stage tilt required for FIB-view imaging which, in turn, is dependent on the stage tilt for lamella fine milling. For waffle milling, FIB-view images were recorded at a stage tilt of 20° in lamella milling orientation, requiring trench milling steps III-IV at –6° stage tilt. The required stage tilt for trench milling steps III-IV (α_Trench milling_) as a function of the stage tilt for FIB-view imaging (α_FIB-view_) can be calculated according to this equation:

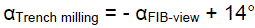

This equation is only valid for the standard geometry used in FIB/SEM instruments employing a shuttle with a pre-tilt of 45°. Care must be taken that the difference between trench milling and FIB-view imaging is not only the stage tilt but also the stage rotation. Trench milling is performed with a stage rotation of 180° and FIB-view imaging with a stage rotation of 0° (Figure 1 – figure supplement 3).

First, the lamella front and the target area were identified in the reflected light channel (excitation wavelength = 470 nm ± 10 nm, no emission filter) and the cellular autofluorescence channel (excitation wavelength = 470 nm ± 10 nm, emission wavelength = 515 nm ± 15 nm), respectively and z-stack recorded (pixel size = 168 nm, 20 z-slices, z-step = 1 µm) using the Odemis software (Delmic). To target the junction between cells, it was sufficient to roughly measure its distance from the sample surface in the Odemis software. For smaller targets, the lamella front can be protected by a layer of metal organic platinum using the GIS and reference marker milled as described (Capitanio *et al*., in preparation)

### Fine milling of waffle lamellae

The surfaces of the block after trench milling were coated with a thick layer of metal organic platinum using the GIS in trench milling orientation and in lamella milling orientation for 15 s and 3 times 20 s, respectively. Fine milling was performed at a stage tilt of 20° in lamella milling orientation (Figure 1 – figure supplement 3c). First, all material below the target area was removed at an ion current of 3 nA. Next, regular cross-sections were used to thin the lamella to an xy-size of 30×10 µm with a beam current of 3 nA and then down to 30×6 µm with 1 nA (Figure 2 – figure supplement 2a). At this point, a notch similar to^21^, but larger, was introduced at current of 0.5 nA (Figure 2 – figure supplement 2b). Subsequently, the width of the milling patterns was reduced in x to 18-20 µm and the lamella thinned at 0.5 nA to 3 µm, 0.3 nA to 1.5 µm and 0.1 nA to 0.8 µm using regular cross-section patterns (Figure 2 – figure supplement 2c). Fine milling was performed with rectangle patterns milled in parallel, aiming for a lamella thickness of 250 nm at 50 pA. Finally, the stage was tilted to 19° and 20.5-21°, removing material only on bottom and top, respectively, at 30 pA.

**Figure 2 – figure supplement 2.**
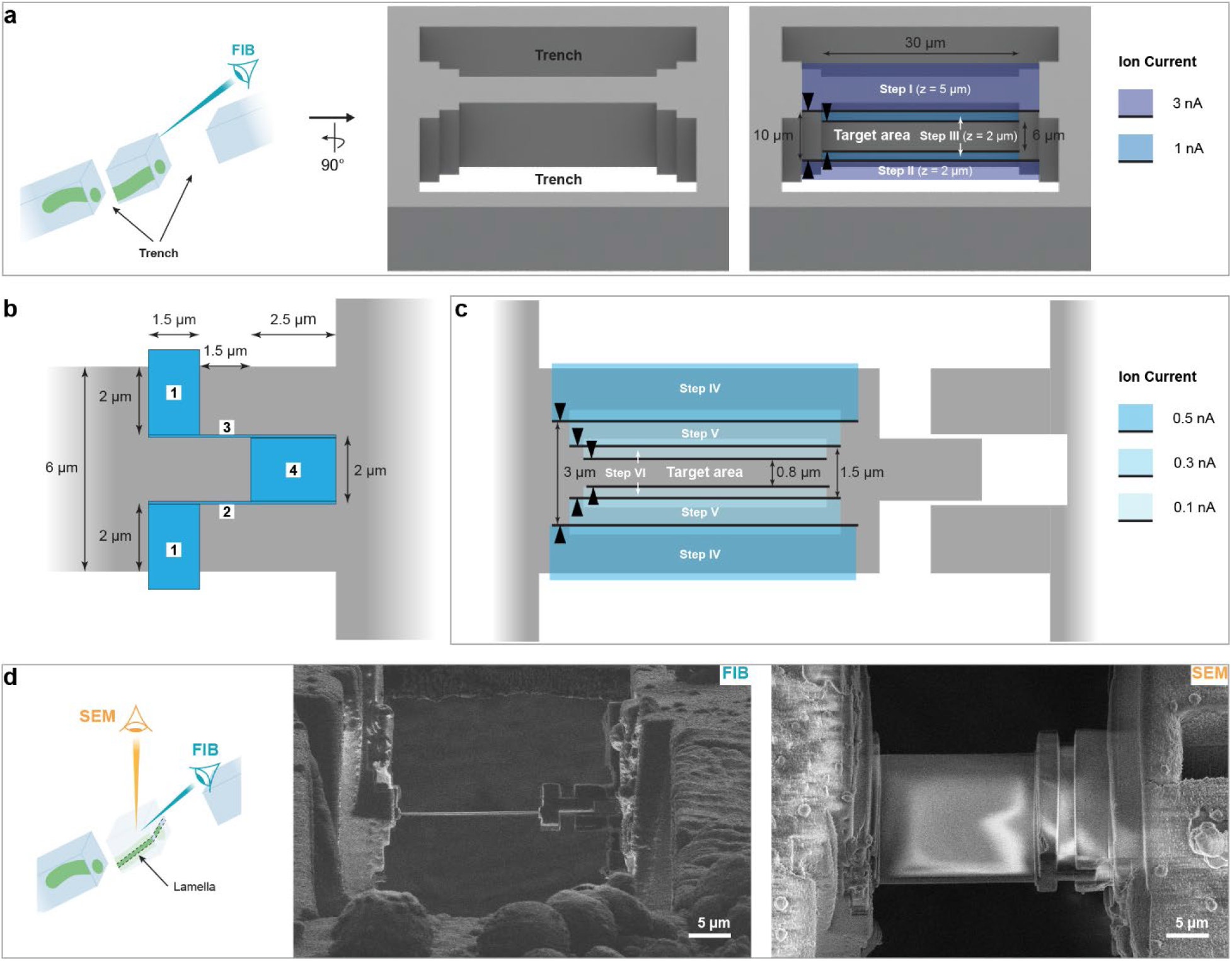
Fine milling of waffle lamellae. **a**-**c**, Scheme of the milling patterns used for waffle lamella preparation. The block after trench milling and rotating the stage into the lamella milling orientation is shown in (**a**). First the lamella is thinned to a thickness of 6 µm. Then a broad notch is milled (**b**) and the lamella further thinned down (**c**). All thinning steps in (**a**) and **(c)** were milled with regular cross-section with milling direction towards the target area indicated by black arrowheads. Fine milling was then performed with rectangle patterns milled in parallel. **d**, FIB and SEM image of a waffle lamella.

### Serial Lift-Out

The EasyLift system (Thermo Fisher Scientific) installed on a Scios and an Aquilos 2 was used for the lift-out experiments. Prior to the lift-out, a copper adapter was attached to the needle of the EasyLift system as described in^22^.

### Lift-out

The first step of the lift-out experiment was targeted trench milling, as described for waffle milling in the previous sections. The width of the block between the trenches depends on the size of the target area. The trenches, however, should be roughly 10 µm wider than the desired width of the final lift-out block as it must be detached from the bulk sample on both sides.

We use redeposition milling to attach a copper adapter to the needle of the micromanipulator, the lift-out block to the copper adapter and the sections to the receiver grid. The pattern used for redeposition milling is an array of regular cross-section with lateral dimensions of 0.5 x 2.5 µm at a distance of 0.25 µm with the multi-pass option set to one. The cross-section patterns are placed along the whole interface of the surfaces that are being attached, on the copper face and the 2.5 µm long dimension is always perpendicular to the attached surface. The milling direction is always away from the interface and the z-dimension of all cross-sections set to 4 µm. On FIB/SEM instruments with imperfect ion beam optics alignment, it can be beneficial to make increase the size of the cross-sections and the distance between them by 40 – 60 % in all directions. A more detailed description of the milling patterns and the whole workflow can be found in the step-by-step protocol (Pöge *et al*., protocol was accepted for publication).

We used two distinct lift-out orientations: (i) from the trench milling orientation, the lift-out direction is roughly in the plane of the sample surface or (ii) from the lamella milling orientation where it is roughly perpendicular to the sample surface (Figure 1 – figure supplement 3d). For (i) the copper adapter was positioned in the center of the block-face directly after trench milling (Figure 3 – figure supplement 1b). Afterwards, the lift-out was performed as described in^22^. For (ii), after trench milling the stage was moved to the lamella milling orientation and a thick layer of metal organic platinum applied by opening the GIS for 3×20 s. Next, all material above and below the target area was removed at 3 nA and the top of the block was polished at 1 nA. Then, the copper adapter was positioned in the center on top of the lift-out block and redeposition milling was used to attach it to the lift-out block (Figure 3 – figure supplement 1a). Only then was the lift-out block detached from the bulk sample on both sides and subsequently extracted.

**Figure 3 – figure supplement 1.**
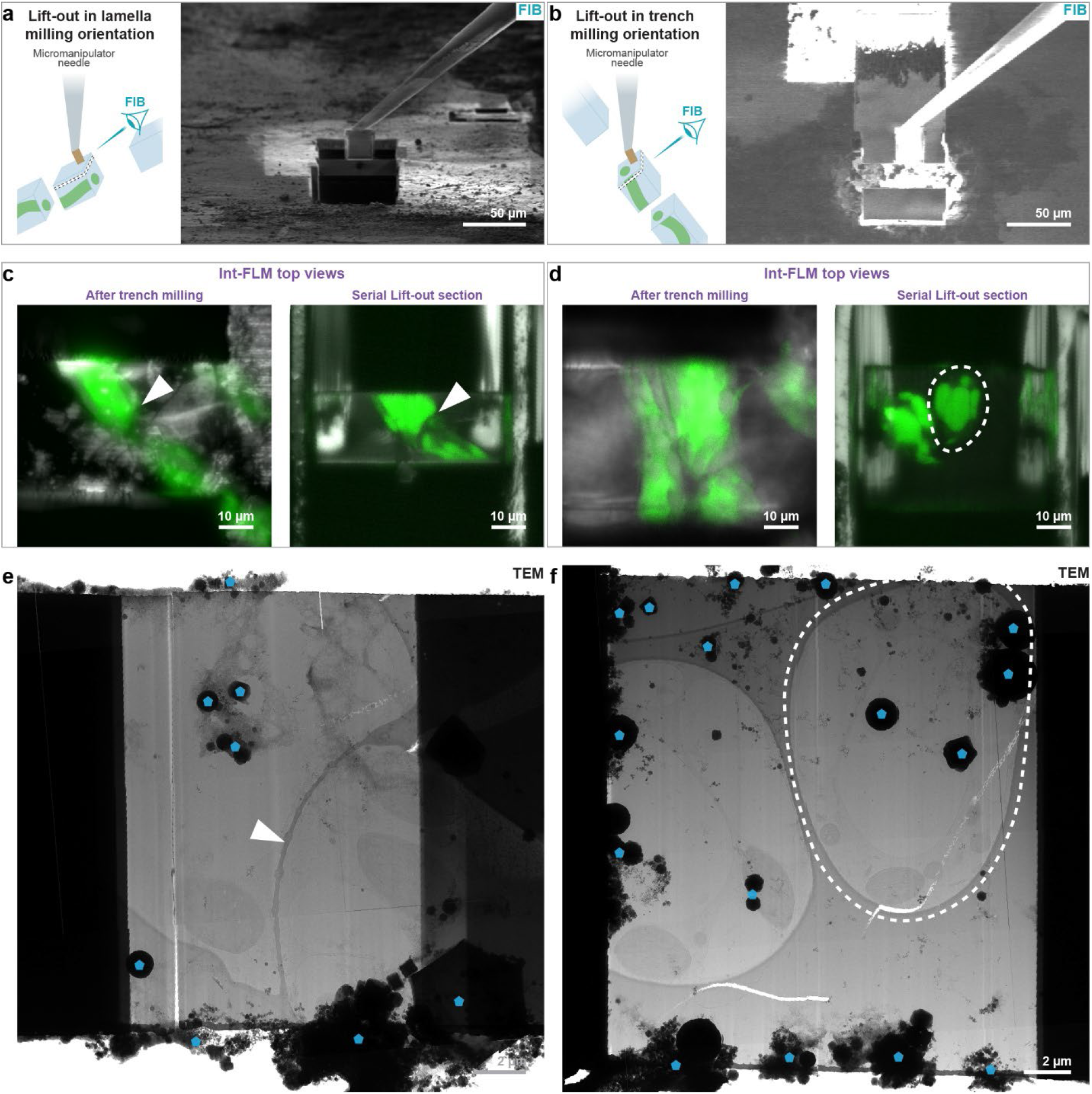
Lift-out in different stage orientations on *P. patens* protonemata. The left panel shows data for the lift-out with the stage in lamella milling orientation, the right panel for the lift-out in trench milling orientation. **a**,**b,** FIB images with the needle attached to the lift-out block. **c**,**d**, Int-FLM top views. The left image and the right image were recorded after trench milling and after sectioning, respectively (reflected light in gray and cellular autofluorescence in green). **e**,**f**, TEM overviews of the final lamellae obtained from the sections in (**c**) and (**d**). The outlines of protonemata cross-sections are marked by dashed lines and cell junctions by white arrowheads. In **(e)** and **(f)** some ice crystals contaminating the lamella surface which were introduced during transfers between microscopes are marked by blue transparent pentagons.

Lift-out from different stage orientations can help imaging cells with preferred orientation on the grid from different directions. Lift-out in trench milling orientation results in cross-sections of protonemata if the axis of the filament runs parallel to the lift-out direction (Figure 3 – figure supplement 1c,e).. In contrast lift-out in lamella milling orientation results in longitudinal sections of protonemata (Figure 3 – figure supplement 1d,f).

### Lift-in using single-sided attachment

For the single-sided lift-in attachment, copper 4-pin half-moon grids (Pelco FIB-Lift-Out-Grids, Plano GmbH, Wetzlar, Germany) were used as receiver grids. After loading, the grid was rotated into trench milling orientation and marker lines milled on either side at the tip of each pin. After rotating back to lamella milling orientation, the needle, with the lift-out block attached, was reinserted and the lower front edge of the lift-out block was aligned to the marker line (Figure 3 – figure supplement 2a). Redeposition milling at 1 nA was used to attach the bottom part of the lift-out block to the pin (Figure 3 – figure supplement 2b). The needle was then moved 200 nm up in z to apply some strain. A horizontal line pattern was milled to section the lowest 3-4 µm of the lift-out block and a vertical line pattern above the attachment patterns at the interface between lift-out block and pin used to fully detach the remaining lift-out block. This process of attachment and sectioning was repeated until the lift-out block was sliced up. More details about the exact milling patterns can be found in the step-by-step protocol (Pöge *et al.*, protocol was accepted for publication).

**Figure 3 – figure supplement 2.**
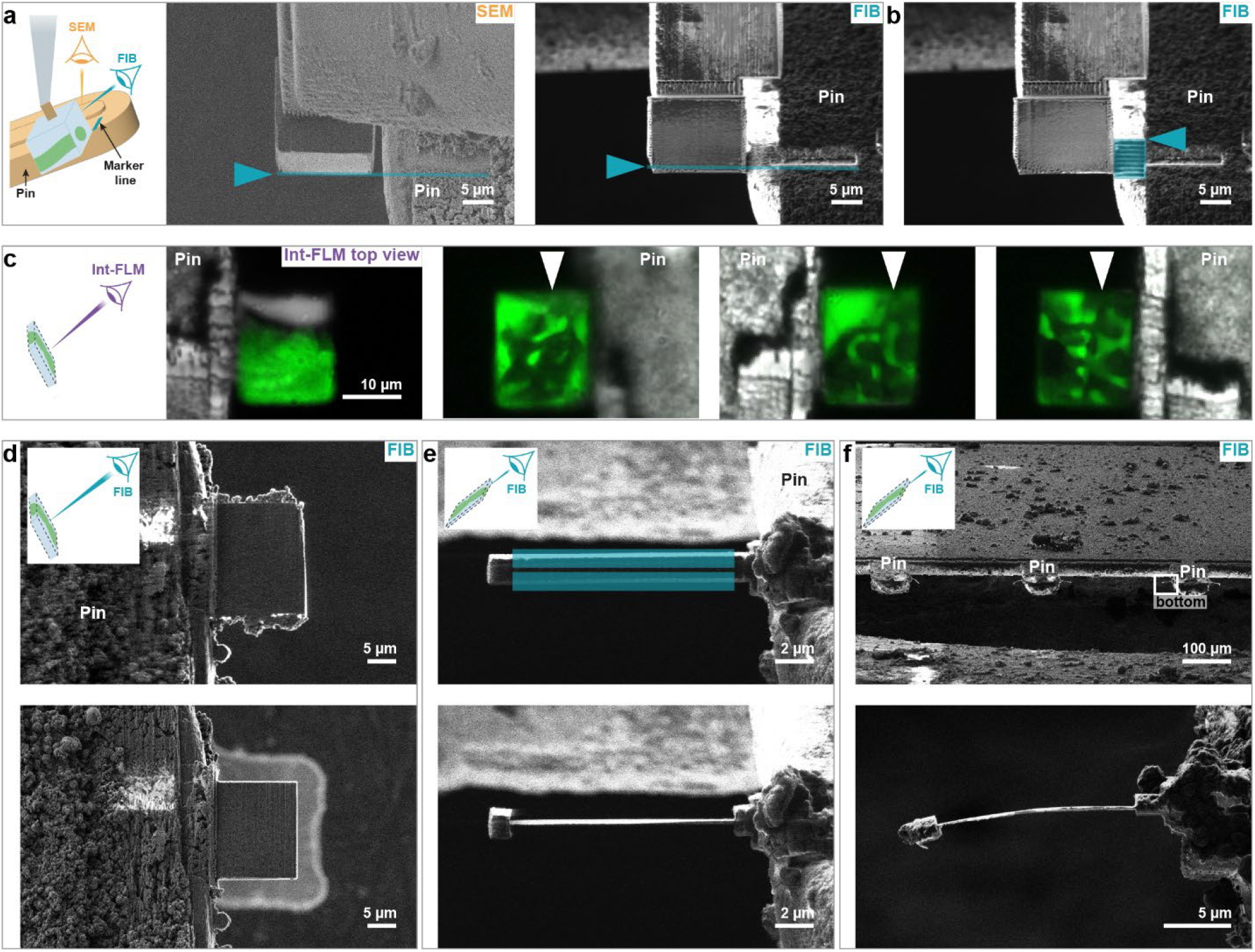
Serial Lift-Out with single-sided attachment of a *P. patens* protonemata. **a**, Alignment of the lift-out block to the marker line (blue line and arrowhead) in FIB and SEM. **b**, Attachment of the lower part of the lift-out block on one side. The patterns for redeposition milling are indicated in blue. **c**, Int-FLM top views of single-sided attached Serial Lift-Out sections acquired in trench milling orientation (reflected light in gray, cellular autofluorescence in green). The junction between cells is only contained the three right-most sections, marked by white arrowheads. **d**, Section trimming in trench milling orientation. A section before and after trimming is shown in the top and bottom panel, respectively. **e**, Single-sided attached lamella before and after fine milling in the top and bottom panel, respectively. The milling patterns are indicated in blue. A 1.5×1.5 µm block is left opposite of the attachment site. **f**, Single-sided attached lamellae often bend during or after fine milling.

### Section trimming

For each section, top views images were recorded with the int-FLM in trench milling orientation to identify sections containing the target area (Figure 3 – figure supplement 2c). Next, the GIS was used for 20 s in trench milling orientation. The section was subsequently trimmed down at 3 nA to primarily contain the target area (Figure 3 – figure supplement 2d), and the front edge on top was polished at 1 nA. To protect the new lamella front, the GIS was used for 15 s and 3×20 s in trench milling and lamella milling orientation, respectively. This trimming step minimizes the size of the lamella and generates a smooth, coated lamella front, which makes the subsequent fine milling faster and easier.

### Fine milling of lamellae with single-sided attachment

The sections were thinned in lamella milling orientation (stage tilt = 15°) at 1 nA to 2.5 µm and at 0.3 nA to 1.5 µm using regular cross-section patterns. Opposite of the attachment site, a 1.5×1.5 µm block was left which prevents folding^53^ of the lamella (Figure 3 – figure supplement 2e). Fine milling was done at 50 pA using rectangle patterns milled in parallel set at a distance of 250 nm from one another, followed by tilting to 14° and 15.5°-16° while milling only on bottom and top, respectively, at 30 pA. Lamellae with single-sided attachment had the tendency to bend (Figure 3 – figure supplement 2f) making fine milling difficult. Additionally, they often broke during transfers into the TEM and displayed sample movement during data acquisition. By reducing the width of the lamella from the attachment site and increasing its depth along the attachment site, this could be reduced but not completely prevented, especially for lamellae thinner than 200 nm. Thus, we strongly recommend using the double-sided attachment procedure as described below.

### Lift-in using double-sided attachment from the sides

For the double-sided attachment, as described in^22^, the lift-out block must fit precisely between two neighboring 400 mesh bars of a 400×100 mesh grid (∼40 µm). As such, the trench milling patterns were adjusted so that the final lift-out block had a width of 42-45 µm. When mounting the 400×100 mesh grid into the shuttle, it is crucial that the 100 mesh bars are aligned horizontally. Marker lines were milled in trench milling orientation in the middle of 400 mesh bars (Figure 3 – figure supplement 3a). Lift-in was performed in lamella milling orientation (stage tilt = 15°). After precise vertical alignment of the 400 mesh bars by rotating the stage, the width of the lift-out block was adjusted to fit precisely between them by FIB milling using cross-sections at 1 nA. The lower front edge of the lift-out block was then aligned to the marker lines of the grid bars in the FIB and SEM channels (Figure 3 – figure supplement 3b).

**Figure 3 – figure supplement 3.**
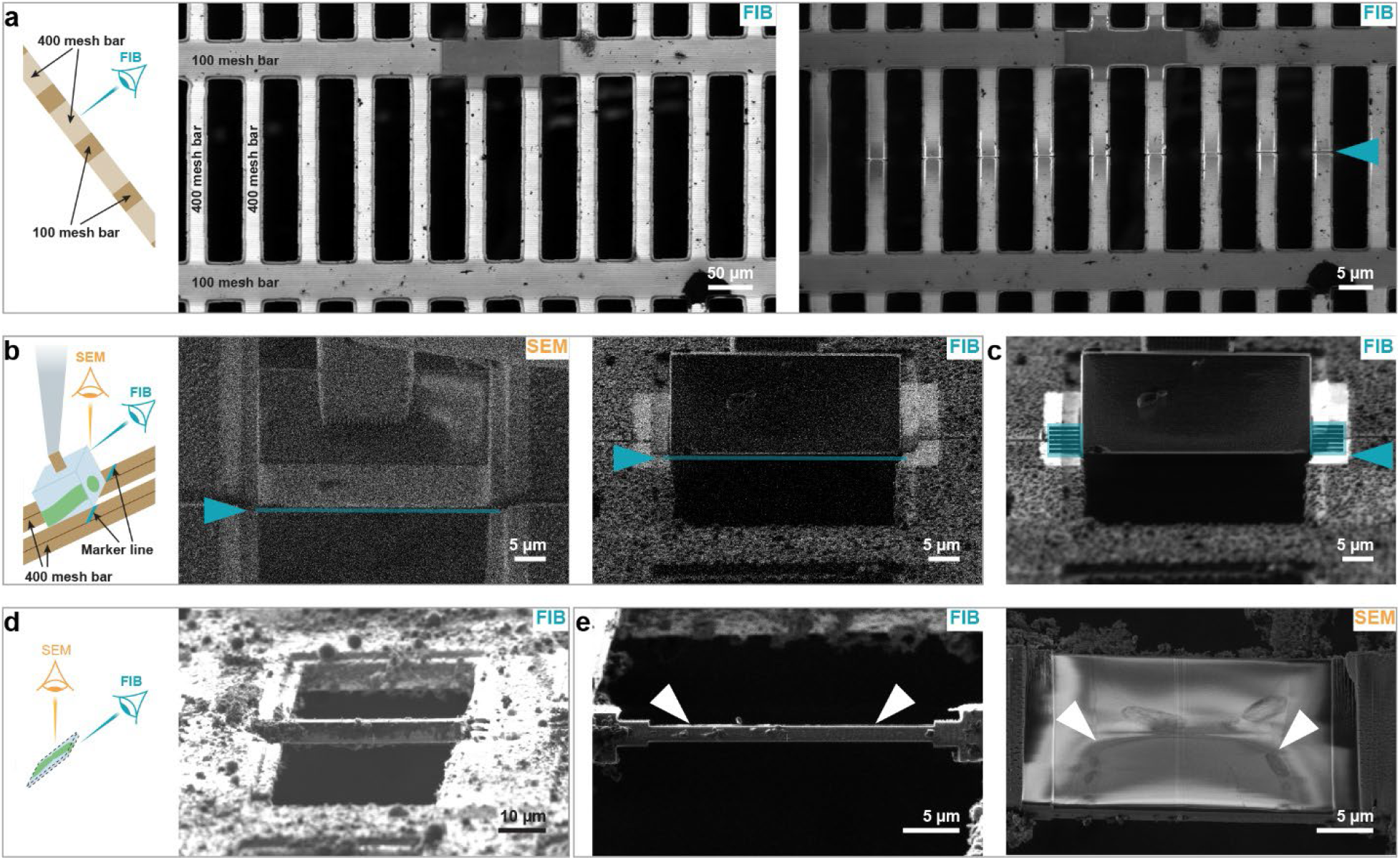
Serial Lift-Out with double-sided attachment from the sides. **a**, Milling of marker lines for the alignment of the lift-out block. Grid bars before and after marker line milling in the left and right panel, respectively. Marker lines are indicated by a blue arrowhead. **b**, Alignment of the lift-out block to the marker lines. SEM and FIB images are shown in the left and right panel, respectively. The bottom front edge of the lift-out block is indicated by a blue line and the marker line by a blue arrowhead. **c**, Attachment of the lift-out block on both sides. Patterns for redeposition milling are indicated in blue. **d**, FIB image of a lift-out section. **e**, FIB and SEM image of the section in the left and right panel, respectively, after the top layer was removed. The SEM image reveals the location of the cell junction as target area (white arrowheads) which was, subsequently, used to minimize the width of the milling patterns for lamella fine milling.

Redeposition milling at 1 nA was used to attach the lower part of the lift-out block to the grid bars on both sides (Figure 3 – figure supplement 3c). Then, the needle was moved up in z by 200 nm. A horizontal line pattern was milled to detach the lower 4 µm of the lift-out block and two vertical line patterns were milled on both sides above the attachment patterns at the interface between lift-out block and grid bars to fully detach the remaining lift-out-block. This process of attachment and sectioning was repeated until the whole lift-out block was sliced up. More details about the exact milling patterns can be found in the step-by-step protocol (Pöge *et al.*, protocol was accepted for publication).

Similar to single sided attachment, after sectioning, top views were recorded for each lamella precursor with the int-FLM in trench milling orientation. Regions that did not contain the target area were removed at 3 nA and the front edge on top was polished at 1 nA (Figure 4). To protect the new lamella front, the GIS was used for 15 s and 3×20 s in trench milling in lamella milling orientation, respectively.

Fine milling was performed in lamella milling orientation (stage tilt = 15°) in several steps. First, the top of the section was removed at 1 nA. Subsequently, the lateral location of the target area was determined based on either the int-FLM top views recorded prior to section trimming or an SEM block-face image (Figure 3 – figure supplement 3e). Using this information, the milling patterns were centered around the target area to reduce the width of the lamella to a minimum. Afterwards, sections were thinned at 1 nA to 2.5 µm, 0.3 nA to 1.5 µm and 0.1 nA to 0.8 µm using regular cross-section patterns. Fine milling was done at 50 pA using rectangle patterns with a distance of 250 nm milled in parallel, followed by tilting to 14° and 15.5°-16° while milling only on bottom and top, respectively, at 30 pA.

### Lift-in using double-sided attachment from below

The idea of the attachment from below is to mill two plateaus into neighboring grid bars, both at the same height and parallel to the bottom of the lift-out block. Thus, the lift-out block can be placed on the plateaus instead of being placed between grid bars. Before the lift-out experiment, a 400×100 mesh grid was loaded into the FIB at room temperature and modified. In trench milling orientation, with an ion current of 64 nA, 8×90 µm wide (xy-size) rectangular cavities were milled into the neighboring 400 mesh bars with the lower end of the patterns in the middle of rectangle (Figure 3 – figure supplement 4a). The modified grids can be stored until further use.

For the lift-out experiment, a modified 400×100 mesh grid was loaded with the 100 mesh bars aligned horizontally in lamella milling orientation (stage tilt 15°). Before the first Serial Lift-Out sections could be attached, the actual plateaus were milled in the back of the cavities (Figure 3 – figure supplement 4b). First, regular cross-section patterns were milled, preparing the rough plateaus, followed by two rectangle patterns milled in parallel on either side to assure that the plateaus have the same height. The width of the resulting attachment site is between 50-55 µm. The lift-out block must be 1-2 µm slimmer to fit in the gap but must still overlap several micrometers with the plateaus on both sides. This was achieved by adjusting the width of lift-out block during trench milling.

After the lift-out, the bottom of the lift-out block was polished resulting in a surface parallel to the plateaus. Next, the center of the lift-out block was aligned to the front edge of the plateaus (Figure 3 – figure supplement 4c). Redeposition milling was used to attach the lower part of the lift-out block on both sides to the plateaus (Figure 3 – figure supplement 4d). After the attachment, the needle was moved up in z by 200 nm and a line pattern was used to detach the lowest 3 µm of the lift-out block. This process of attachment and sectioning was repeated until the whole lift-out block was sliced up. Afterwards, top views were recorded with the int-FLM for each section followed by trimming and fine milling as described in the previous section.

**Figure 3 – figure supplement 4.**
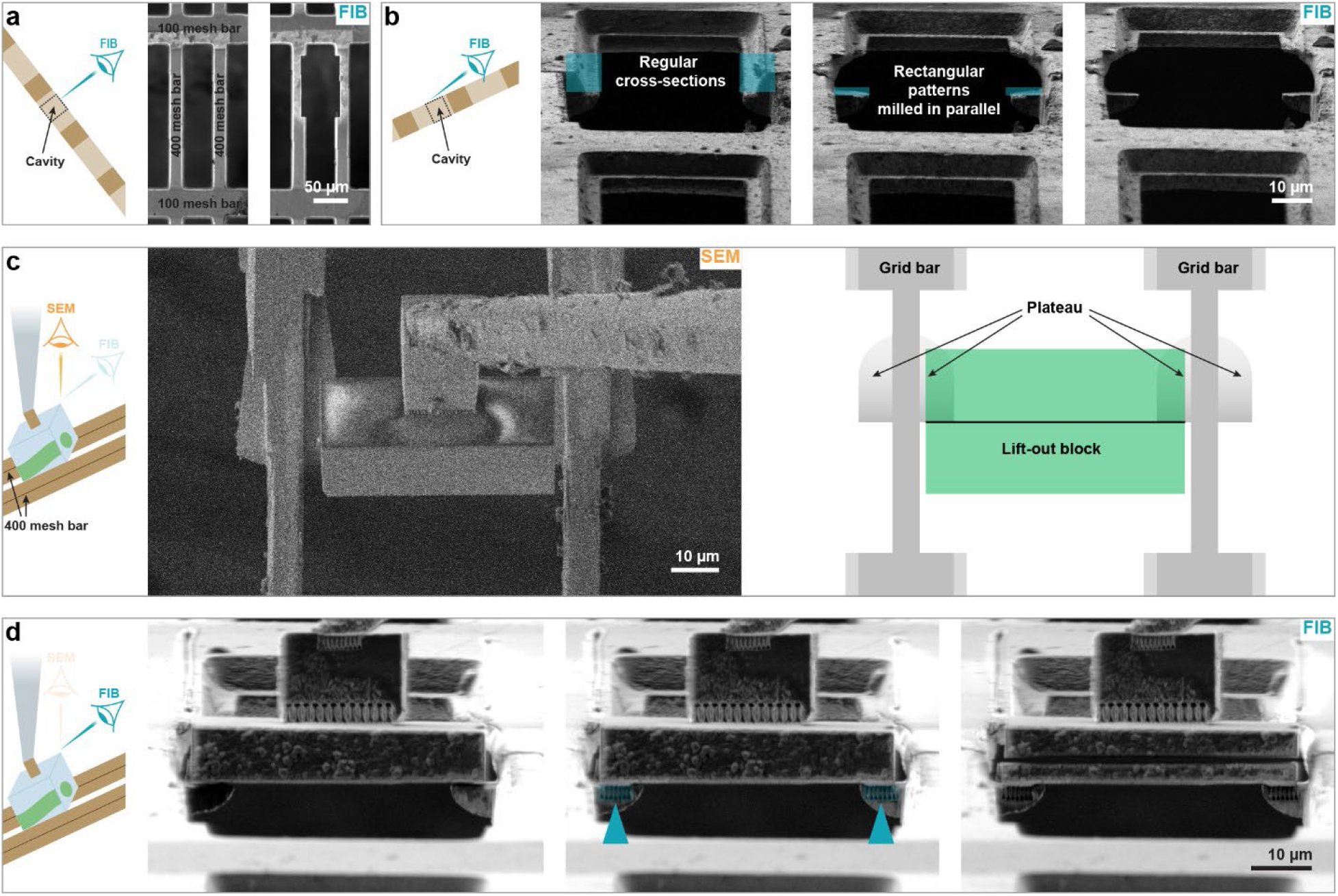
Serial Lift-Out with double-sided attachment from below. **a**, Preparation of 8×90 µm cavities in trench milling orientation. **b**, Preparation of the plateaus in lamella milling orientation. Left panel: FIB image of cavities before milling. Middle panel: after milling regular cross-section patterns (xyz: 9 x 8 x 20 µm) generating the rough shape of the plateaus. Due to stage drift, they can have different heights. Right panel: After rectangle patterns were milled in parallel (xyz: 9 x 1 x 20 µm) to even out height differences. This allows the attachment of the bottom of the lift-out block to both plateaus from below. Milling patterns are indicated by blue rectangles. **c**, Alignment of the lift-out block with respect to the plateaus in the SEM. The center of the block should be roughly on top of the front edge of the plateaus (sketch in right panel). Precision of the placement, however, is not imperative. **d**, the lift-out block placed on the plateaus, after attachment from below and after sectioning in the left, middle and right panel, respectively. Patterns for redeposition milling are depicted in blue.

### In-carrier high-pressure freezing for *P. patens*

Commercially available 3mm Type A and Type B carriers (Leica Microsystems, Wetzlar, Germany) were used for in-carrier HPF. Type A has a 100 µm deep well on one side and a 200 µm well on the other. Type B has only a single 300 µm deep well and is flat on the other side. Carriers were incubated overnight in a 1:1 mixture of 4 M NaOH and commercial bleach, rinsed 3× with water, 1x with Acetone and dried. Subsequently, the type only B carriers were coated by dipping them into cetyl palmitate solution (0.5 (m/v) in diethyl ether), removing excess of liquid with filter paper and drying carriers with the flat side facing up on filter paper. As freezing buffer, a 1:1 mixture of growth medium and 40% Ficoll 400 was used. The 100 µm well of the type A carrier was filled with 2 µL freezing buffer and the droplet spread over the entire well with the pipet tip. A small patch of *P. patens* tissue was placed in the droplet, air bubbles were removed with tweezers and the excess of buffer removed with filter paper until the meniscus of the droplet almost vanished. The type B carrier was placed on top with the flat side down, the sandwich squeezed together and immediately high-pressure frozen. Afterwards, the sample containing type A carriers were recovered and stored until further use.

**Figure 3 – figure supplement 5.**
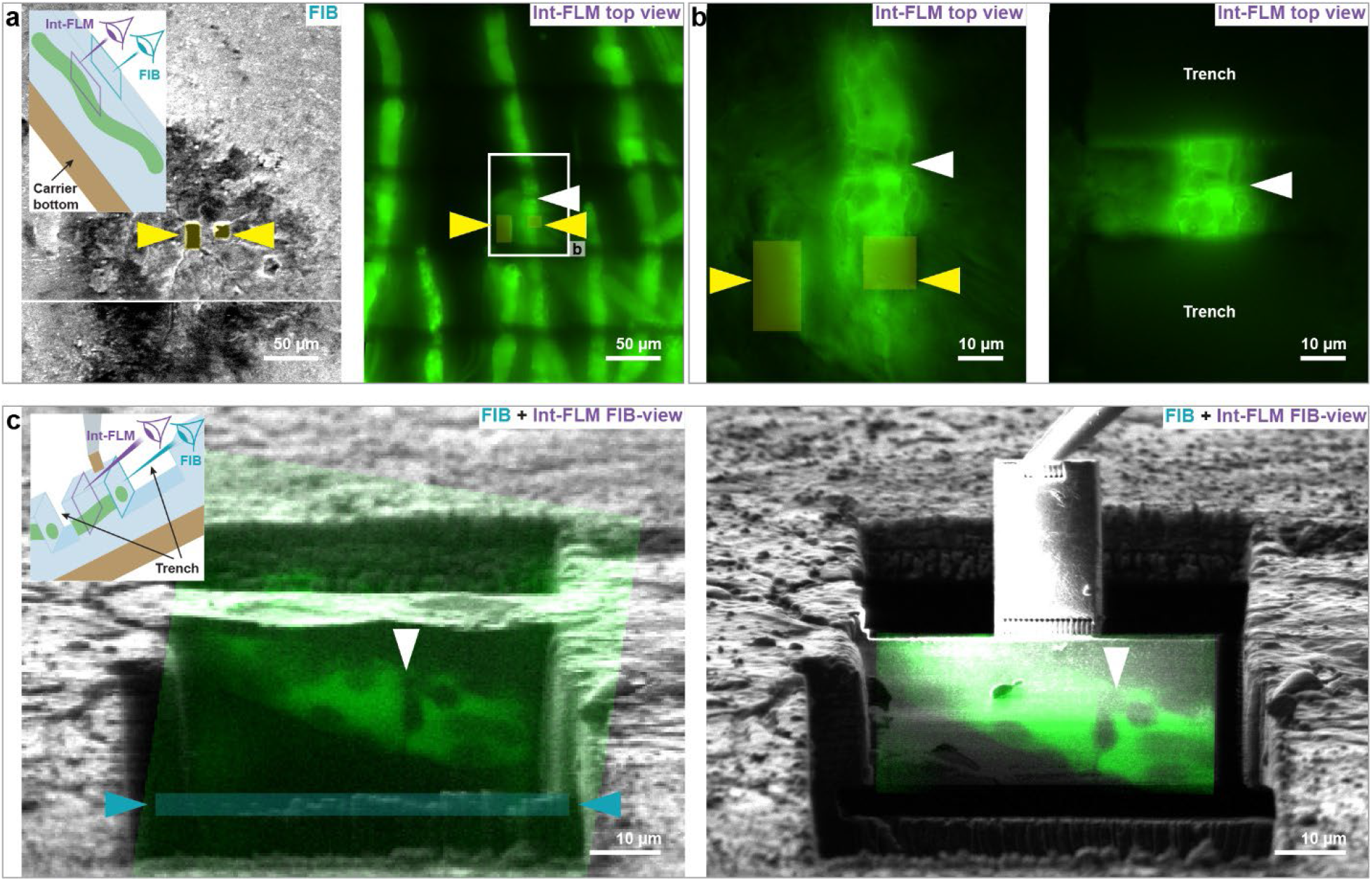
Preparation of in-carrier frozen HPF samples for lift-out (*P. patens* protonemata). **a**, Screening of in-carrier frozen HPF sample using the int-FLM. FIB and a montage of int-FLM top view images in left and right panel, respectively. The field-of-view in (**b**) is marked by a white rectangle. **b**, Int-FLM images before and after trench milling in the left and right panel, respectively. In (**a**) and (**b**), marker patterns are labelled with yellow arrowheads and yellow rectangles. **c**, Milling of the undercut, aided by the int-FLM for in-carrier frozen samples. Overlay of FIB images (gray) and int-FLM FIB-view image of cellular autofluorescence (green) before and after milling the undercut in the left and right panel, respectively. Both images were recorded in lamella milling orientation. The undercut is marked by a blue rectangle. In (a)-(c) cell junctions are labelled by white arrowheads.

In-carrier HPF samples were not screened in our FLM as they don not fit into the commercial shuttle. Hence, they were screened using the int-FLM. After loading the samples into the FIB, the sample surface was protected by a layer of metal organic platinum for 45 s by using the GIS with the stage in trench milling orientation. Afterwards, marker patterns were milled (xy-dimensions: 6 x 16 µm and 8 x 8 µm at a distance of ∼10 µm with z-dimension = 0.5 µm) and top view images were recorded with the int-FLM, covering area of 1000 µm x 1000 µm around the marker pattern (Figure 3 – figure supplement 5a). Targeted trench milling was performed as described for waffle grids (Figure 3 – figure supplement 5b) with the exception that target areas were also selected based on their proximity to the sample surface, which could be roughly estimated based on where the fluorescence was localized within the acquired int-FLM z-stacks.

To fully detach the lift-out block from an in-carrier HPF sample, an additional undercut is necessary. First, the axial location of the target area was determined either by FIB-view imaging with the int-FLM (Figure 3 – figure supplement 5c) or SEM block-face imaging (Figure 2d). With the stage in lamella milling orientation, the front of the lift-out block protected by metal organic platinum and the undercut milled below the target area at 3 nA. Afterwards, the lift-out was performed as described in the previous section either from the trench milling or the lamella milling orientation.

### Lamella sputter coating after fine milling

Lamellae were sputter coated after fine milling. The sputtering systems on the used Aquilos 1 and Aquilos 2 introduce 5-15 nm sized spheroidal platinum particles onto the lamella surfaces. The sputtering conditions were set to U = 1 kV, p = 0.1-0.2 mbar, I = 10-15 mA, t = 2-4 s. These particles aid frame and tilt-series alignment during TEM data processing. The fiducial based tilt-series alignment resulted in higher quality tomograms than patch tracking approaches.

### Waffle workflow for *P. patens* phyllids

The waffle grids of *P. patens* phyllids were prepared similar to protonemata. A cleaned and cetyl palmitate coated 6 mm type B carrier was placed with the flat side facing up on filter paper and a 3 µL droplet of freezing buffer placed on it. Next, a 50 mesh EM-grid with continuous support film (Carbon coated Formvar, Graticules Optics Ltd, Tonbridge, UK) was placed on the droplet with the support side facing down and excess of buffer removed until the grid was resting flat on the carrier. A 3 µL droplet of freezing buffer was added on top of the grid and distributed to fill all wells formed by support film and grid bars and air bubbles removed with sharp tweezers. A small piece of phyllid was cut from the gametophore with fine surgical tools and placed in the center of the grid. A second coated 6 mm type B carrier was placed with the flat side facing down on top, the sandwich squeezed together and excess of buffer removed with a filter paper. Next the sandwich was mounted for HPF and squeezed tightly for 10 s right before freezing. Areas of tissue that were damaged in the process could be identified in the FLM (Figure 5 – figure supplement 1a-c). In areas with largely intact cells, individual chloroplasts can be resolved (Figure 5 – figure supplement 1b), whereas for damaged cells, the autofluorescence signal is diffusely distributed throughout the whole cell (Figure 5 – figure supplement 1c). Lamellae were otherwise prepared by waffle milling or lift-out as described in the previous sections.

**Figure 5 – figure supplement 1.**
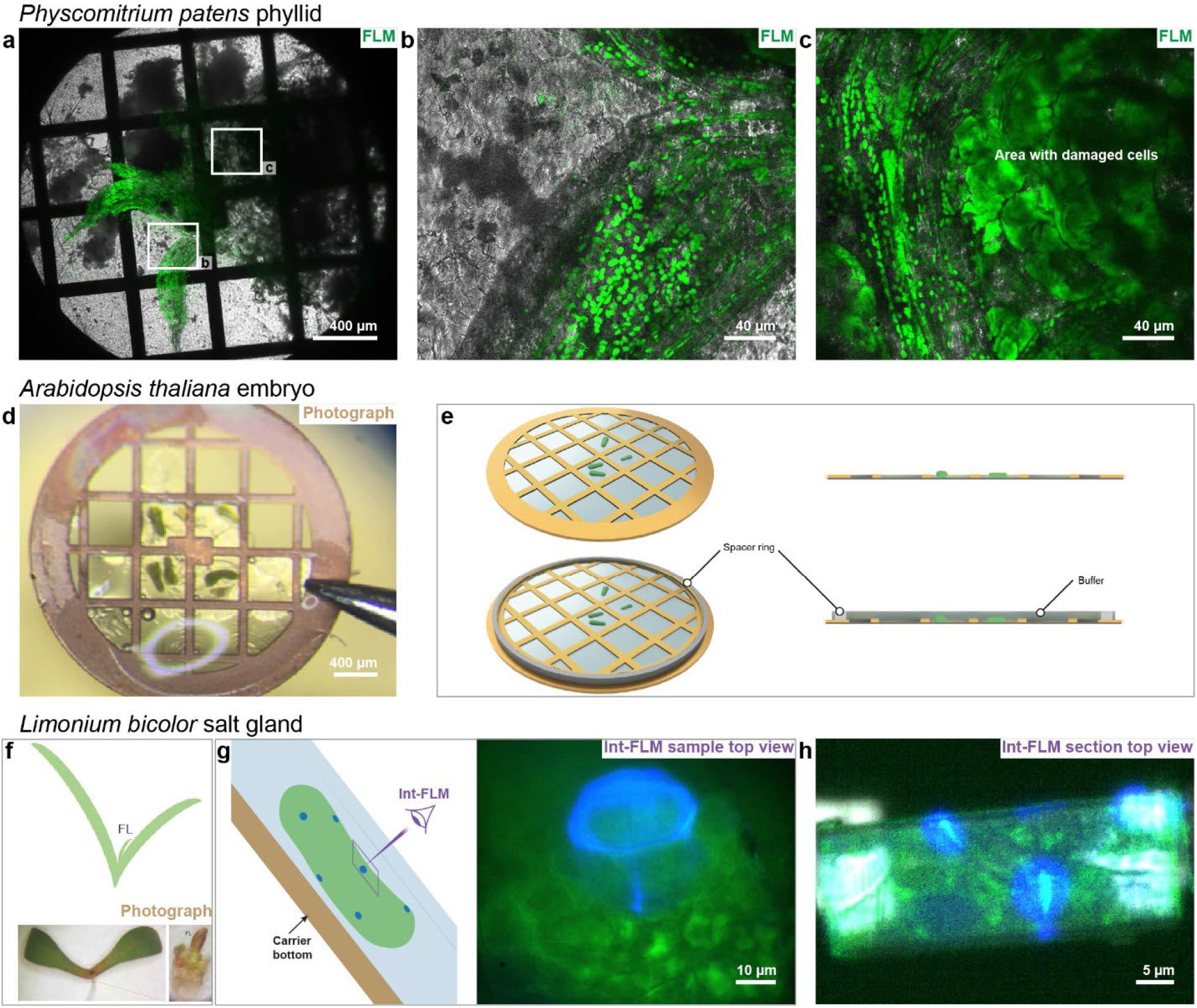
HPF workflows applied to other tissues and species. **a**-**c**, *Physcomitrium patens* phyllids. **a**, FLM overview of a phyllid waffle grid. **b**, Grid square containing seemingly intact tissue. **c**, Grid square containing damaged tissue. **d**,**e** *Arabidopsis thaliana* embryos. **d**, Image of embryos placed in the squares of a 50 mesh grid. **e**, Schematic of a waffle grid with and without a spacer ring. Embryos fit into grid squares but are thicker than the grid bars and would therefore be damaged during freezing (top panel). The addition of a spacer ring, as described in^21^, prevents squeezing damage and can allow high-pressure freezing of thicker samples. **f**-**h**, Salt gland of *Limonium bicolor*. **f**, Dissection of the first true leaf (FL). The top panel shows a scheme of the FL location in 2 weeks old plants. The bottom panel are photographs of a whole two weeks old plant before and the dissected FL in the left and right panel, respectively. **g**, int-FLM top view image of the salt gland of in-carrier frozen FL. **h**, int-FLM image top view image of a Serial Lift-Out section containing a fraction of the salt gland. In (**g**) and (**h**) autofluorescence of the salt gland is shown in blue, the cellular autofluorescence in green and the reflected light in gray.

### The waffle workflow for extracted *Arabidopsis thaliana* embryos

*A. thaliana* ecotype Columbia (Col_0) was grown 5-7 weeks in soil at 22.5°C with a photoperiod of 16/8 h (day/night_)._ The isolation of embryos was performed on the preparation table of the Leica EM-ICE high-pressure freezer using its integrated binocular microscope. As freezing buffer, 20% Ficoll 400 (w/v) dissolved in PBS was used. The first eight siliques were cut with fine scissors and placed on a glass slide in a drop of freezing buffer. Next, embryos were hand dissected with sharp tweezers as described in reference^54^ and collected. A 50 mesh EM-grid with continuous support film (Carbon coated Formvar, Graticules Optics Ltd, Tonbridge, UK) was placed with the support facing down in a 3 µm droplet of freezing buffer on the flat side of a cetyl palmitate coated 6 mm Type B HPF carrier. The excess of buffer was blotted away with filter paper until the support film was flat on the carrier. Several embryos were picked up with tweezers and placed in the wells formed by grid bars and support film (Figure 5 – figure supplement 1d). A spacer ring (Provider) with a height of 50 µm was added onto the grid and filled with a 2 µL drop of freezing buffer. Air bubbles were removed with tweezers. Afterwards, the sandwich was closed with a second 6 mm type B carrier flat side facing down, the sample high-pressure frozen, the grid recovered and stored in a grid box in LN_2_.

### Targeting the salt gland of in-carrier frozen *Limonium bicolor*

The culture conditions *L. bicolor* Kuntze, fluorescence microscopy detection of salt glands and HPF procedures are similar to those described previously^55^ with minor modifications. Namely, *L. bicolor* seeds were stored in a refrigerator at 4°C before use. The seeds were then surface disinfected in 70% (v/v) ethanol for 5 min and subsequently in 6% (w/v) sodium hypochlorite (freshly prepared) for 20 min and then thoroughly washed with deionized water. Fully filled seeds were selected and planted in MS culture medium plates. These plates were incubated in a plant tissue culture room at a temperature of 26/22°C (day/night), a photoperiod of 16/8 h (day/night) and a light intensity of 600 μmol·m-2·s-1. After 2 weeks, the first true leaf (FL, Figure 5 – figure supplement 1f) of the whole plant was dissected with an thin knife, placed in the 300 µm deep cavity of a 3 mm Type B HPF carrier filled with with 20% (w/v) Ficoll 400. Afterwards, the cavity was closed off with a second Type B carrier, flat side facing down, the specimen was high-pressure frozen and the sample containing carrier recovered. Fluorescence targeting of the salt gland was performed using the int-FLM recording cellular autofluorescene (excitation wavelength = 470 nm ± 10 nm, emission wavelength = 515 nm ± 15 nm), autofluorescence of salt glands (excitation wavelength = 385 nm ± 10 nm, emission wavelength = 432 ± 18 nm nm) as well as the reflected light (excitation wavelength = 385 nm ± 10 nm, no emission filter, Figure 5 – figure supplement 1g). Serial Lift-Out with the attachment from below was then performed as described in the previous sections (Figure 5 – figure supplement 1h).

### TEM Imaging

One part of the TEM data was collected on a Titan Krios G2 (Thermo Fisher Scientific) with BioQuantum energy filter (Gatan, Pleasanton, California, USA) and K2 summit camera (Gatan). Automated data acquisition was performed in SerialEM^56,57^ (version 4.1). Lamella montages were recorded at 8700x magnification (pixel size = 2.6 nm). Tomograms were acquired at 42000x, 53000x or 64000x magnification (pixel sizes of 3.52 Å, 2.62 Å and 1.98 Å, respectively), at target defoci of –4 µm to –6 µm, using a dose symmetric tilt scheme between ±50°to ±60° with an angular increment of 2° or 3°, taking the pretilt of the lamellae into account. The exposure time was kept constant for all tilt images resulting in total doses per tilt-series of 80-125 e^-^/Å^2^. Typically, 10 frames were recorded per tilt image and saved in TIF-file format.

The other part of the data was recorded on a Titan Krios G4 equipped with a Selectris X energy filter and a Falcon 4i camera (Thermo Fisher Scientific). Lamella montages were recorded at a magnification of 8700x (pixel size = 2.8 nm). Tomograms were recorded at 42000x or 64000x (pixel sizes of 2.93 Å and 1.89 Å, respectively) with defoci between –3 µm and –6 µm. Data was acquired using the Tomography 5 software package version 5.12.0 (Thermo Fisher Scientific) and stored in EER file format.

## Data preprocessing

Tilt-series were processed using the TOMOMAN^58^ (version 0.6.9) pipeline. Movie frames were motion corrected in MotionCor2^59^ (version 1.4.7). EER frames were grouped to obtain between 8 and 20 frames per movie. Bad tilt images, i.e. improperly tracked images or blurry images due to lamella instabilities, were removed after manual inspection and dose-weighting was performed according to^60^ using the corresponding TOMOMAN scripts. If tilt-series contained at least 5 platinum fiducials introduced by sputtering evenly distributed over the field-of-view, IMOD^61^ (version 4.11.25) fiducial tracking was employed for tilt-series alignment. Otherwise, AreTomo^62^ version 1.3.3 was used for tilt-series alignment. Afterwards, 4x binned tomograms were reconstructed with the weighted back projection algorithm implemented in IMOD.

For denoising, tilt series containing only the information of odd or even frames were obtained during motion correction. The odd and even tilt-series were preprocessed to generate 4x binned odd and even tomograms using the same alignment files as for the full tilt-series. Denoising was performed in cryo-CARE^63^. First, 800-1000 training volumes with a box size 72^3^ voxel were cropped from the odd and even tomograms. A network was trained for each tomogram over 150 epochs with 75 steps and a Unet depth of 3. The network was then used to denoise the 4x binned tomograms. All TEM data, including tomograms and lamella overviews, were visualized with IMOD^64^ (version 4.11.25). Only slices of denoised tomograms are shown in this work.

## Subtomogram averaging of rubisco

The detailed workflow for subtomogram averaging and spatial analysis for rubisco is described in the Supplementary Information. In brief, a subset of 10 tomograms were recorded on *P. patens* protonemata chloroplasts with defoci between –2.5 and –5 µm. Tilt-series were aligned and reconstructed in IMOD and imported into WARP^65^ (version 1.0.9). Membranes were segmented using MemBrain-seg^66^ and the chloroplast volumes were manually annotated in Amira (Amira 3D 2021.2, Thermo Fisher Scientific) to create a mask for score maps as output of template matching. A first template was created by picking 500 rubisco complexes manually (50 complexes per tomogram) and aligning them in RELION^67^ (version 3.0.5) rubisco was picked by template matching in StopGap^68^ (version 0.7.0). After several rounds of subtomogram alignment and classification in RELION and tilt-series alignment in *M*^69^ (version 1.0.9) the final density map was calculated using 6,180 particles and applying C4 symmetry (Figure 4c). An octamer of large rubisco subunits (monomer UniProt accession P34915) was predicted in AlphaFold3^39^ and docked into the final subtomogram average using the “fit” function in ChimeraX^70^ (version 1.6.1) without further refinement of the model (Figure 4c-d). The spatial analysis of rubisco particles was performed based on 55,000 particles after a first round of classification in RELION similar to^71^. The 3D rendered image of chloroplast membranes and rubisco positions was displayed in ChimeraX using the ArtiaX plug-in^72^.

## Supplementary information

### Supplementary Movies

Video 1: Lift-out in lamella milling orientation on a waffle grid. This approach resulted in longitudinal sections of protonemata as shown in left panel of Figure 3 – figure supplement 1,e.

Video 2: Lift-out in trench milling orientation on a waffle grid. This approach can resulted in cross-sections of protonemata as shown in right panel of Figure 3 – figure supplement 1d,f.

Video 3: Lift-in with single-sided attachment. At this point we had serious recontamination problems in our FIB-SEM instrument. In this case, the marker line was simply milled into the contamination that accumulated on the pin over time. An example for a single-sided attached lamella is shown in Figure 5a,b.

Video 4: Lift-in with double-sided-attachment from the side.

Video 5: Lift-in with double-sided-attachment from below. An example for a lamella with double-sided attachment from below is shown in Figure 3a-c.

Video 6: *Physcomitrium patens* protonema cell. Example of a tilt-series with the corresponding, denoised tomogram. A single slice of the tomogram is shown in Figure 3d.

Video 7: Lift-out in lamella milling orientation on an in-carrier frozen sample.

Video 8: Lift-out in trench milling orientation on an in-carrier frozen sample.

Video 9: *Physcomitrium patens* protonema chloroplast. Example of a tilt-series with the corresponding, denoised tomogram. A single slice of the tomogram is shown in Figure 4a.

Video 10: *Physcomitrium patens* phyllid. Example of a tilt-series with the corresponding, denoised tomogram. A single slice of the tomogram is shown in Figure 5b.

Video 11: *Arabidopsis thaliana* embryo. Example of a tilt-series with the corresponding, denoised tomogram. A single slice of the tomogram is shown in Figure 5d.

Video 12: *Limonium bicolor* salt gland. Example of a tilt-series with the corresponding, denoised tomogram. A single slice of the tomogram is shown in Figure 5f.

### Rubisco subtomogram averaging

A dataset of 10 tomgrams was recorded on chloroplasts (unbinned pixel size = 0.19 nm, defocus range = [-2.5:0.5:-4.5], tilt-range = [-51°:2°:51°] taking the lamella pre-tilt into account), preprocessed in TOMOMAN^1^ (version 0.6.9) and reconstructed in IMOD^2^ (version 4.11.25)using fiducials on the surface of lamellae for tilt-series alignment. Aligned tilt-series were imported into WARP^3^ (version 1.0.9) and the CTF estimated (patch size = 512 pixel, fit range = [3.5-0.8 nm]).

#### Generation of the initial template

500 rubisco particles were manually selected (50 particles in each tomogram), 4x binned subtomograms (voxel size 0.76 nm, box size = 32, particle diameter = 18 nm for all processing steps) were extracted in WARP, the initial Euler angles randomized and aligned in RELION^4^ (version 3.0.5, Refine3D).

#### Template Matching

The resulting average was rescaled for template matching on 8x binned tomograms (voxel size = 1.51nm) in STOPGAP^5^ (version 0.7.0). The reference was lowpass filtered to 4 nm and a smoothed sphere mask with a diameter of 21 nm applied. The score maps were masked as follows: The Chloroplast volume was manually annotated in Amira (Amira 3D 2021.2, Thermo Fisher Scientific) to pick particles only within the lamella and within chloroplasts. Chloroplast membranes were segmented with MemBrain-seg^6^ (using the pretrained model: MemBrain_seg_v10_alpha.ckpt) and grown 3 nm (two 8x binned voxels) to avoid picking particles too close to membranes. Scores within 20 voxels from the edge of the map were excluded to avoid picking particles at the edge of the tomogram (Supplementary Figure 1a). Initially, the 12 000 highest scoring hits (score > 0.25) were selected.

**Supplementary Figure 1.**
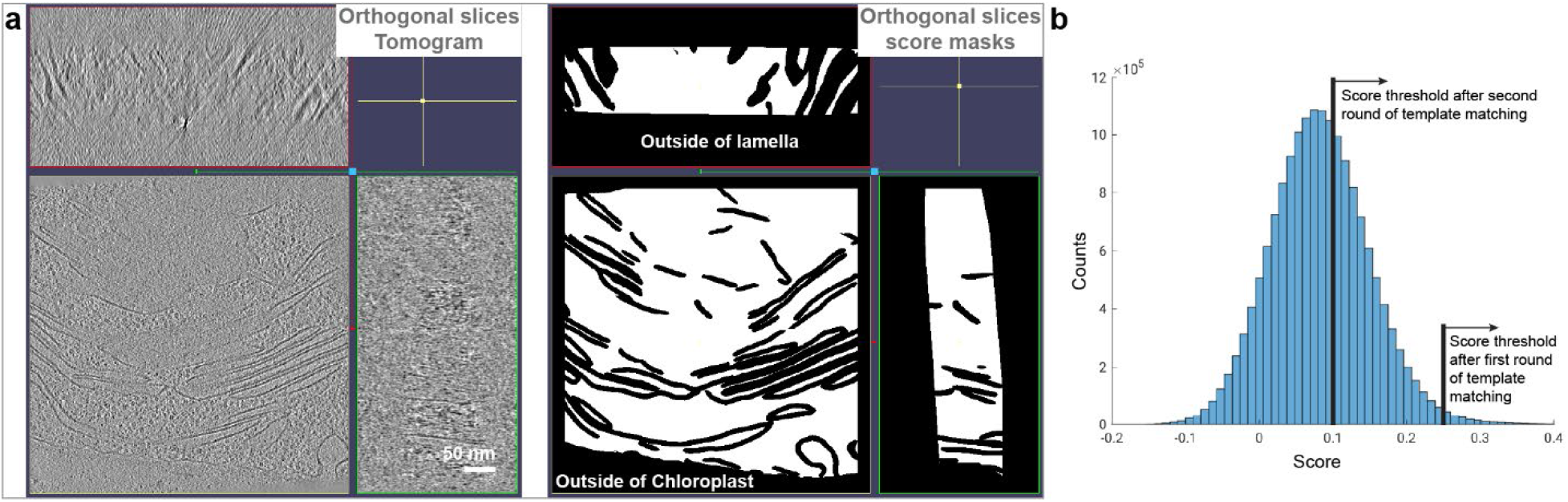
Rubisco template matching. **a**, The mask applied to the template matching score maps. Orthogonal slices of a chloroplast tomogram and the mask applied to the respective score map are shown in the left and right panel, respectively. Particles were only selected in the white area of the mask. **b**, Histogram of the scores inside the mask in **(a).** After the first round of template matching with a poor-quality reference, subtomograms were only extracted at positions with scores higher than 0.25. For the second round, the score threshold was set to 0.1 to avoid false negatives.

#### First round of template matching to create a better template

4x binned subtomograms (voxel size 0.76 nm, box size = 32) were extracted in WARP and all Euler angles randomized. Subtomograms were aligned (Refine3D) and classified (Class3D, 15 classes, regularization parameter = 0.1) in RELION using a sphere mask with a diameter of 18 nm. A simple sphere mask was always used for alignment and classification if not otherwise indicated. Subtomograms showing the expected features and symmetry of rubisco were selected and aligned separately (Refine3D). The resulting average was rescaled and used as template for a second round of template matching on 8x binned tomograms with STOPGAP.

**Supplementary Figure 2.**
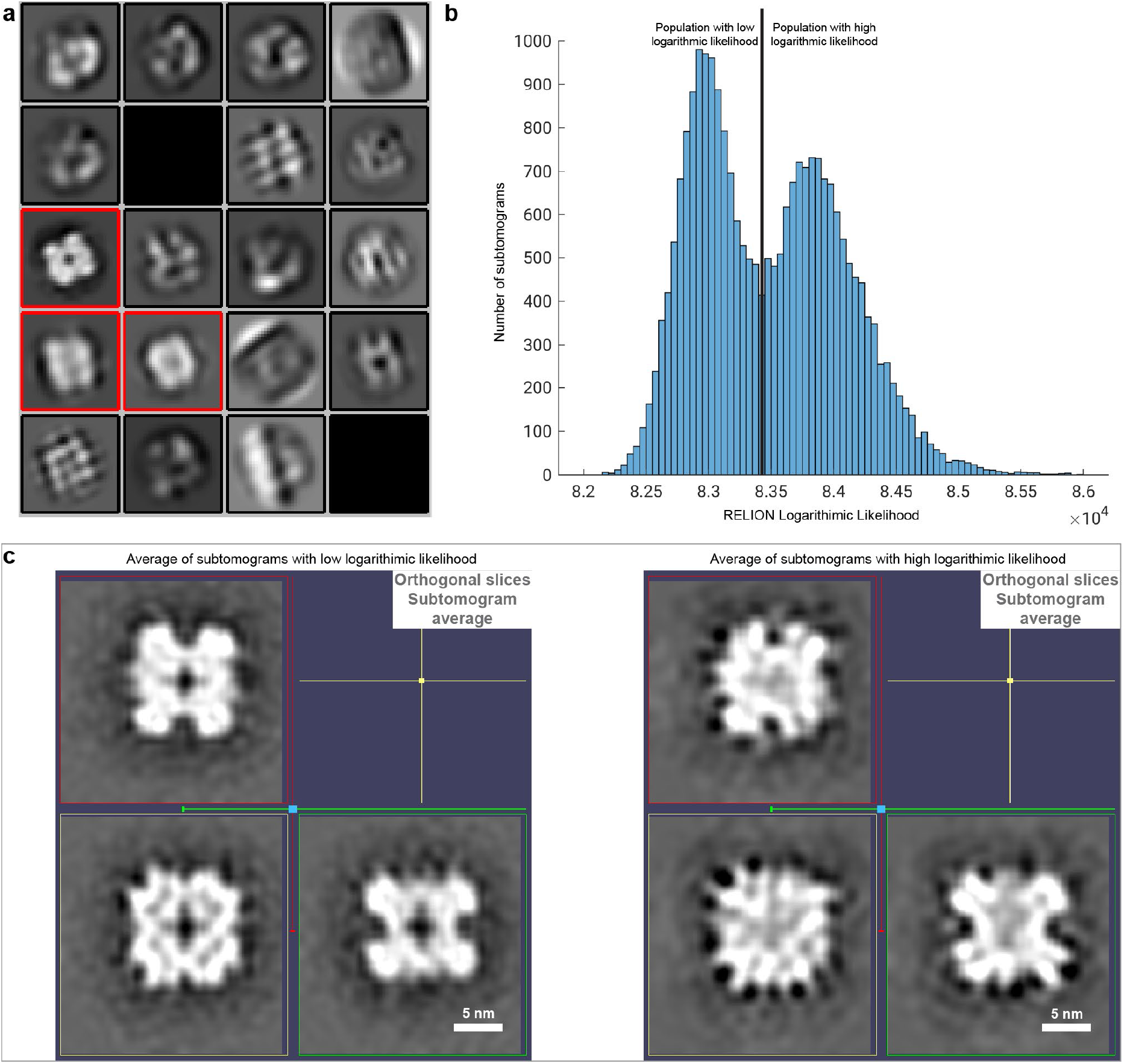
Classification of rubisco subtomograms in RELION. **a**, Class averages as result of classifying a batch of 40 000 subtomograms after the second round of template matching. Class averages that were selected are framed in red. **b,** RELION logarithmic likelihood histogram of subtomograms after tilt-series alignment in M and subtomogram alignment in RELION. Two populations are observed. **c,** The averages after aligning the subtomograms in the population with the low and the high logarithmic likelihood in **(b)** separately are shown in the left and right panel, respectively.

#### Processing of 4x binned subtomograms

To assure that most rubisco particles were picked after the second round of template matching, the threshold for hits was chosen to be less stringent. All particles with a score larger than 0.1 were extracted 4x binned (Supplementary Figure 1b) in WARP resulting in 161 000 subtomograms and the Euler angles were randomized. After an initial alignment (Refine3D), subtomograms were randomly split into 4 batches of 40 000 particles and classified (Class3D, 20 classes, regularization parameter = 0.5, Supplementary Figure 2a). Many class averages showed no similarity with the average used for template matching and the associated subtomograms were excluded from further processing. The remaining 55 000 particles were aligned (Refine3D) using a non-spherical, smoothed mask (MaskCreate). The refined positions after this alignment were used for the calculation of the radial local rubisco density (See below). After an additional round of classification (Class3D, 20 classes, regularization parameter = 0.4), the 24 000 subtomograms assigned to the classes with the highest resolution, were selected, aligned separately (Refine3D) and used for tilt-series alignment in *M*^7^ (version 1.0.9, Refinement of 4x binned data for particle positions, stage tilts, image warp 3×3, volume warp 2×2×1).

#### Processing of 2x binned subtomograms

The 24 000 particles were rextracted as 2x binned subtomograms (voxel size = 0.38 nm, box size = 64 voxel) in WARP. After an initial alignment (Refine3D) with a non-spherical smoothed mask (MaskCreate) the average appeared spiky. The histogram of the logarithmic likelihood as determined by RELION revealed two distinct populations (Supplementary Figure 2b). Processing the 13 000 subtomograms with the higher logarithmic likelihood resulted in a significantly worse average than for the 11 000 subtomograms with lower logarithmic likelihood (Supplementary Figure 2c). Hence, the population with the higher logarithmic likelihood was excluded from further processing.

#### Processing of unbinned subtomograms

So far, the symmetry of rubisco was not considered during subtomogram alignment. According to the TOM toolbox^8^ function “tom_check_symmetry”, the average exhibited C4 symmetry which was applied in all subsequent processing steps. After alignment of the 11 000 subtomograms (Refine3D) with non-spherical smoothed mask and tilt-series alignment in M on 2x binned data, unbinned subtomograms (voxel size = 0.19 nm, box size = 128 voxel) were reextracted in WARP. Subtomograms were aligned (Refine3D), classified (Class3D, 10 classes, regularization parameter = 0.1), 6180 subtomograms in the highest resolved classes selected and separately aligned (Refine3D). The Fourier Schell Correlation (Supplememntary Figure 3a) was calculated based on the half-maps as output of RELION using a smoothed mask calculated on the final average filtered to 1.5 nm, with binarization threshold 0.1 extended by 3 voxels and a soft edge of 15 voxels (MaskCreate). B-factor sharpening was performed in RELION (PostProcessing). The B-factor was estimated to be –429 taking special frequencies below 2 nm into account. The sharpened density map is shown in Supplementary Figure 3b. The flow shard with all processing steps is depicted in Supplementary Figure 4.

**Supplementary Figure 3.**
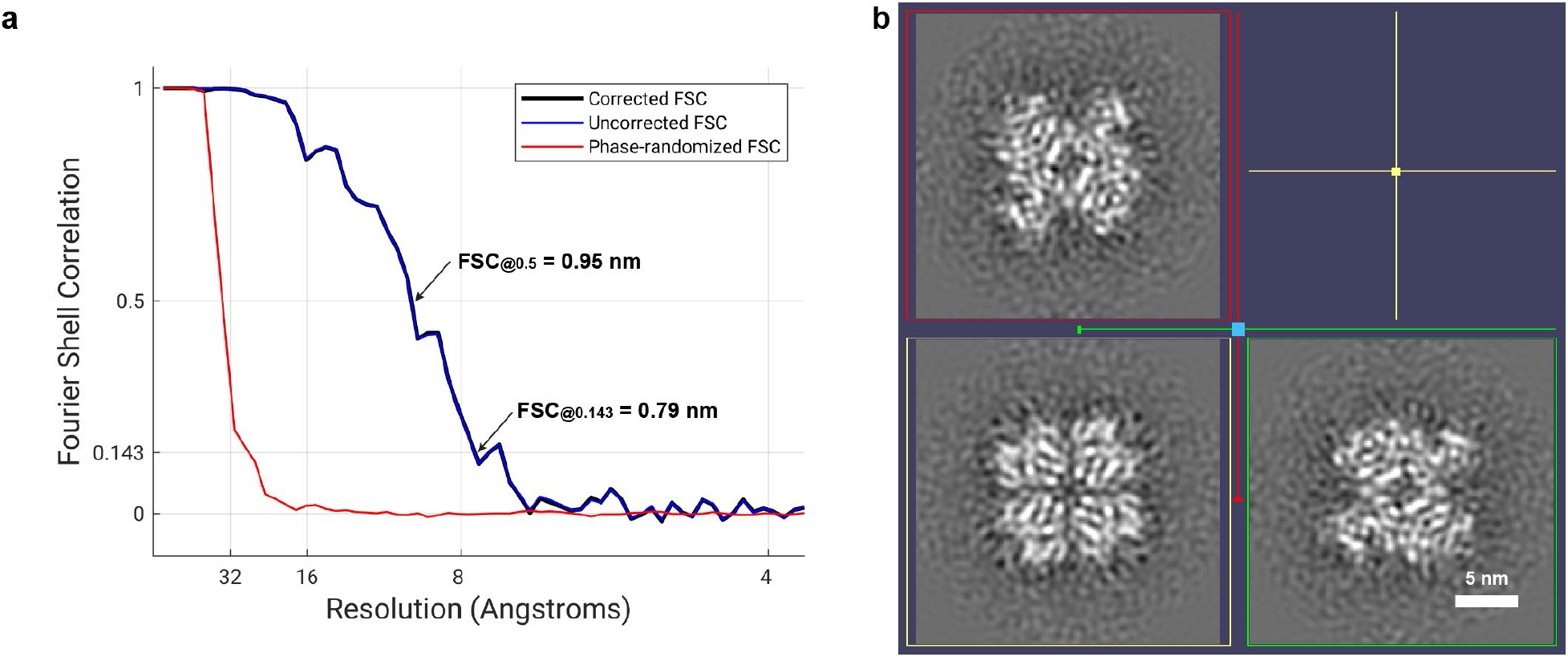
The final rubisco subtomogram average. **a**, Fourier Shell Correlation of the half maps. **b,** Orthogonal slices of the final rubisco subtomogram average which is shown as isosurface representation in Figure 4c.

**Supplementary Figure 4.**
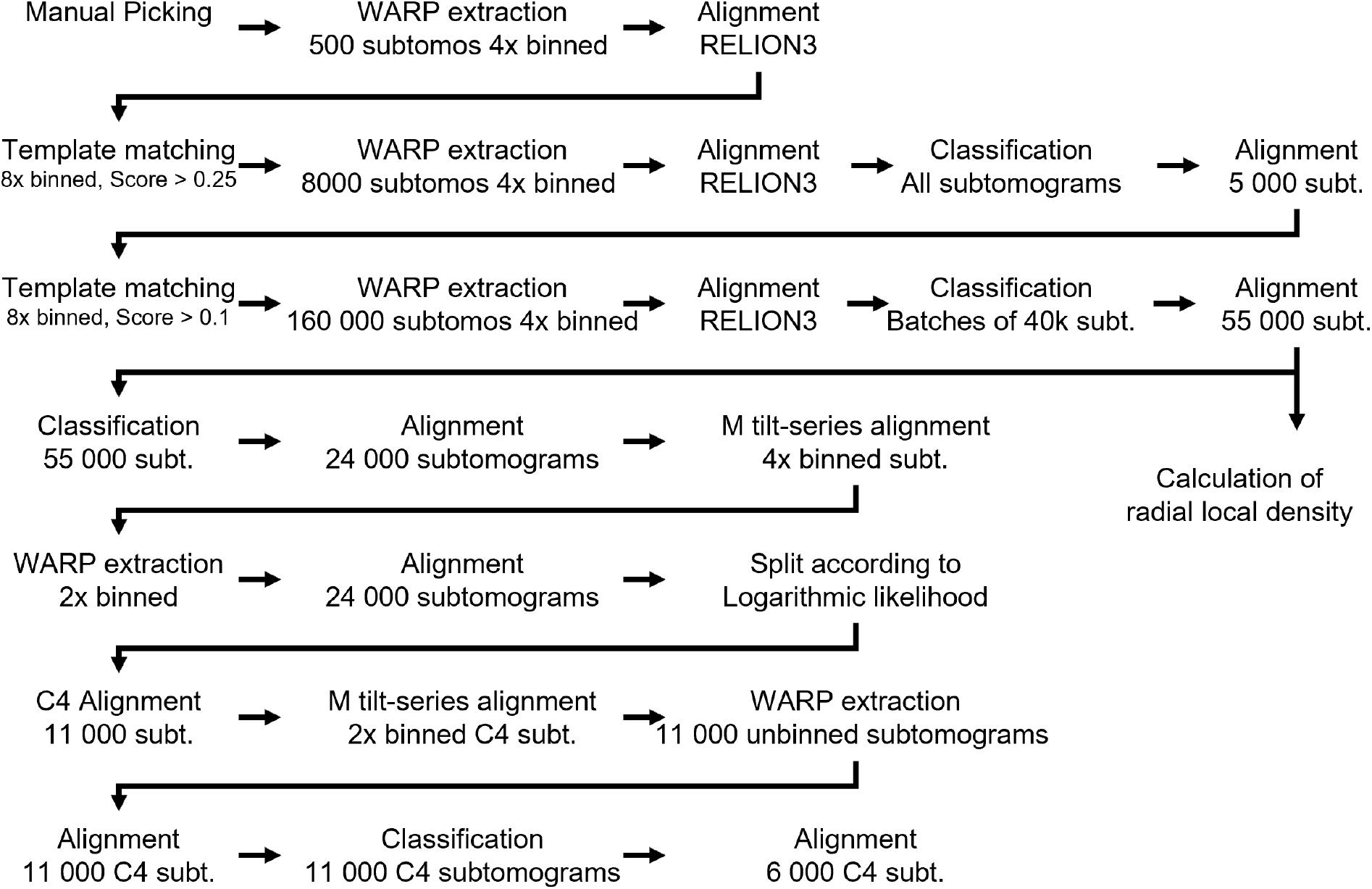
Flow shard of the employed subtomogram averaging wokflow.

### Spatial analysis of rubisco

#### Calculation of the experimental radial local desnity

The radial local density of rubisco was calculated similar to the workflow in^9^ but with a new implementation in MATLAB (MathWorks, version 2022a) and without taking Euler angles into account. The subtomogram coordinates as sum of extraction point plus shift for 55 000 4x binned subtomograms were divided by two, rounded and assigned to the tomogram where they were extracted from. For each tomogram, every rubisco was represented as voxel with value one in a volume of zeros with the same size as the 8x binned tomogram. For every rubisco, the radial local density was calculated as the number of other rubisco complexes found within 50 shells (maximum distance ∼75 nm) with a thickness of 1 voxel (distance increment ∼ 1.5 nm). The number of rubisco complexes per shell was divided by the number of non-zero voxels of the mask which was applied to the score maps (Supplementary Figure 1a) within the same shell. In a last step, this value was divided by the voxel volume to derive the actual rubisco density per shell and average over all complexes resulting in the radial local density for one tomogram. The radial local density for all tomograms separately and the mean ± one standard deviation for all tomograms combined are the green curves in Supplementary Figure 5 and Figure 4d, respectively.

**Supplementary Figure 5.**
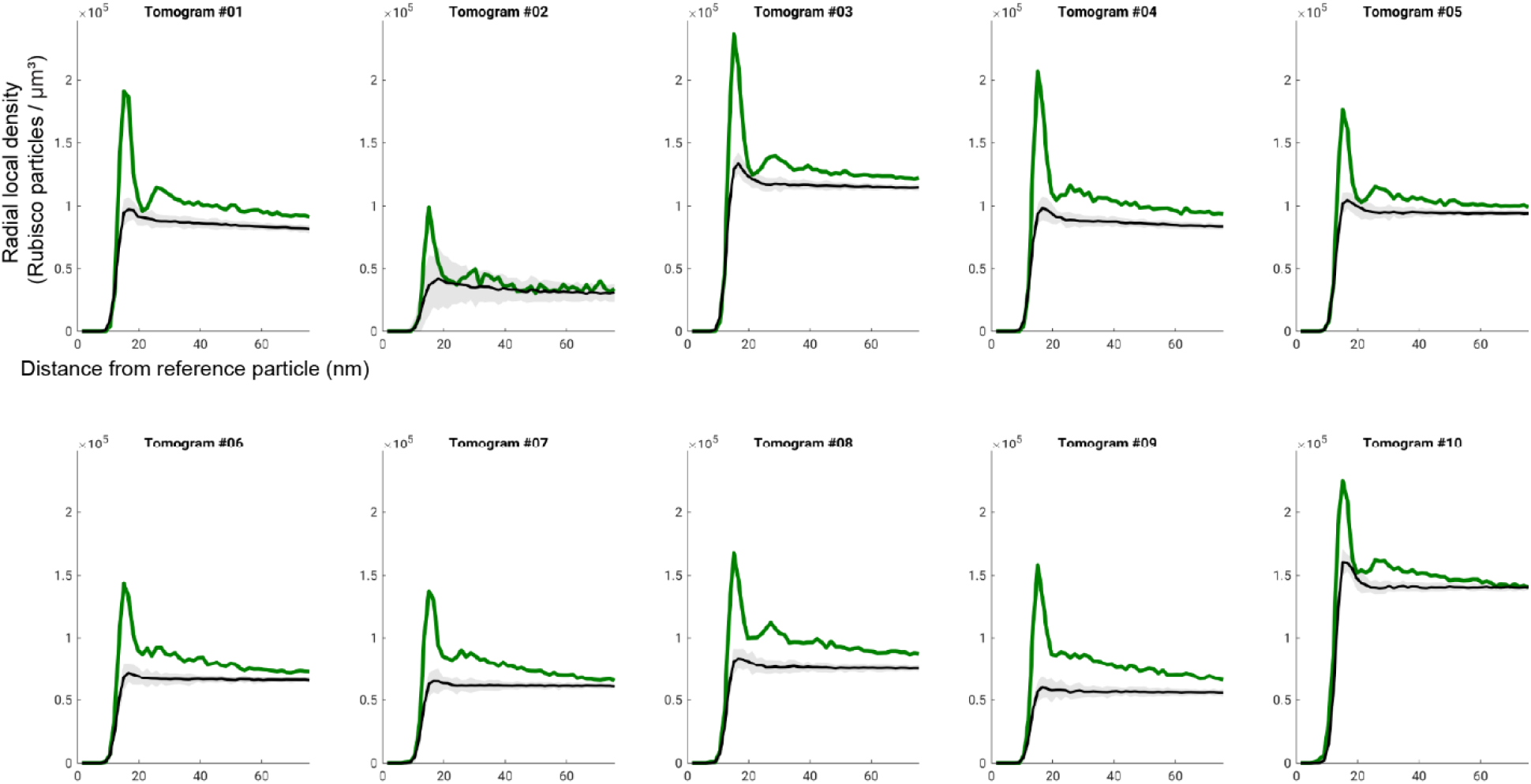
The experimental radial local density of rubisco compared to a simulated random distribution. The experimental radial local density for each tomogram is shown in green, the mean ± three standard deviations for the radial local density of a simulated random distribution in black. For every tomogram, 20 independent simulations were performed.

#### Simulation of a random distribution

For the simulation of the random particle distribution, first the masks (Supplementary Figure 1a) which were applied to the score maps after template matching were modified. All membranes were grown to half the diameter of rubisco (∼8 nm) to assure that random positions are not picked too close to membranes. Next, random voxels in the non-zero region of the mask were picked a particle position until the number of experimentally rubisco complexes was reached. For every randomly selected particle position, the minimum distance to the next neighbor was randomly selected from a normal distribution with a mean of 12 nm (eight 8x binned voxels) and a variance of 1.5 nm (one 8x binned voxel). This was necessary to simulate the initial smooth rise of the radial local density. As the experimentally determined minimum distance between complexes was larger than the diameter of rubisco, particle orientations were neglected, in contrast to a similar analysis of rubisco in the pyrenoid of the green alga Chlamydomonas reinhardtii^10^. With these randomly selected particle positions, the radial local density was calculated as for the experimentally determined rubisco positions. For every tomogram, 20 independent simulations were performed. The mean radial local density ± three standard deviations of the simulated random distribution for all tomograms separately and the mean ± one standard deviation for all tomograms combined are the black curves in Supplementary Figure 5 and Figure 4d, respectively.

#### Differences in experimental and simulated rubisco density

The difference in the radial local density of the experimental data and the simulated random distribution for all tomograms combined (Figure 4d) is not as striking as the difference in for every tomogram separately (Supplementary Figure 5) because the densities vary substantially between different tomograms. This may indicate that some rubisco particles were not picked in some tomograms perhaps due to higher false negative rate in tomograms with lower defocus or with worse quality. Nevertheless, experimental radial local densities for individual tomograms (Supplementary Figure 5) exhibit a peak around 15 nm which is significantly larger than the mean plus three times the standard deviation calculated for the random distribution.

The simulated density is always slightly lower than for the experimental data. Here, the simulate density is systematically underestimated for two reasons: First, not all membranes were segmented by MemBrain-seg, especially if they are at a steep angle with respect to the z-axis of the tomogram (Supplementary Figure 1a). Second, the luminal space between thylakoid membranes is not everywhere zero in the masks applied to score maps. Thus, simulated particle positions were also selected on non-segmented membranes and within the luminal space. As no rubisco positions were found in these regions experimentally, the simulated local rubisco density with the same number particles must be reduced. Irrespective of that systematic problem, we think the peak height in the experimental data is sufficient to suggest a non-random distribution of rubisco within Chloroplasts of the moss *Physcomitrium patens*.

### Success rates of vitrification and lamella preparation

#### High-Pressure Freezing of *P. patens*

For this work, we high-pressure froze 45 *P. patens* samples (Supplementary Figure 6a). From 21 of these samples (47%), we obtained vitreous lamellae. The HPF results are not consistent and sample dependent. Sometimes even lamellae within the same grid square a few micrometers apart would differ in vitrification state.

#### Prepared lamellae vs lamellae imaged in a TEM

In total, 216 lamellae were prepared. Of 74 on-grid milled waffle lamellae 80% survived the transfer into the TEM and could be imaged. For a fraction of the waffle lamellae not all material below the lamella was removed by FIB milling. These “shadow lamellae” prevent cryo-ET data acquisition. For single-sided attachment, double-sided attachment from the side and double--sided attachment from below 31, 49 and 99 lamellae, respectively, were prepared of which 58%, 76% and 90%, respectively, could be imaged (Supplementary Figure 6b).

**Supplementary Figure 6.**
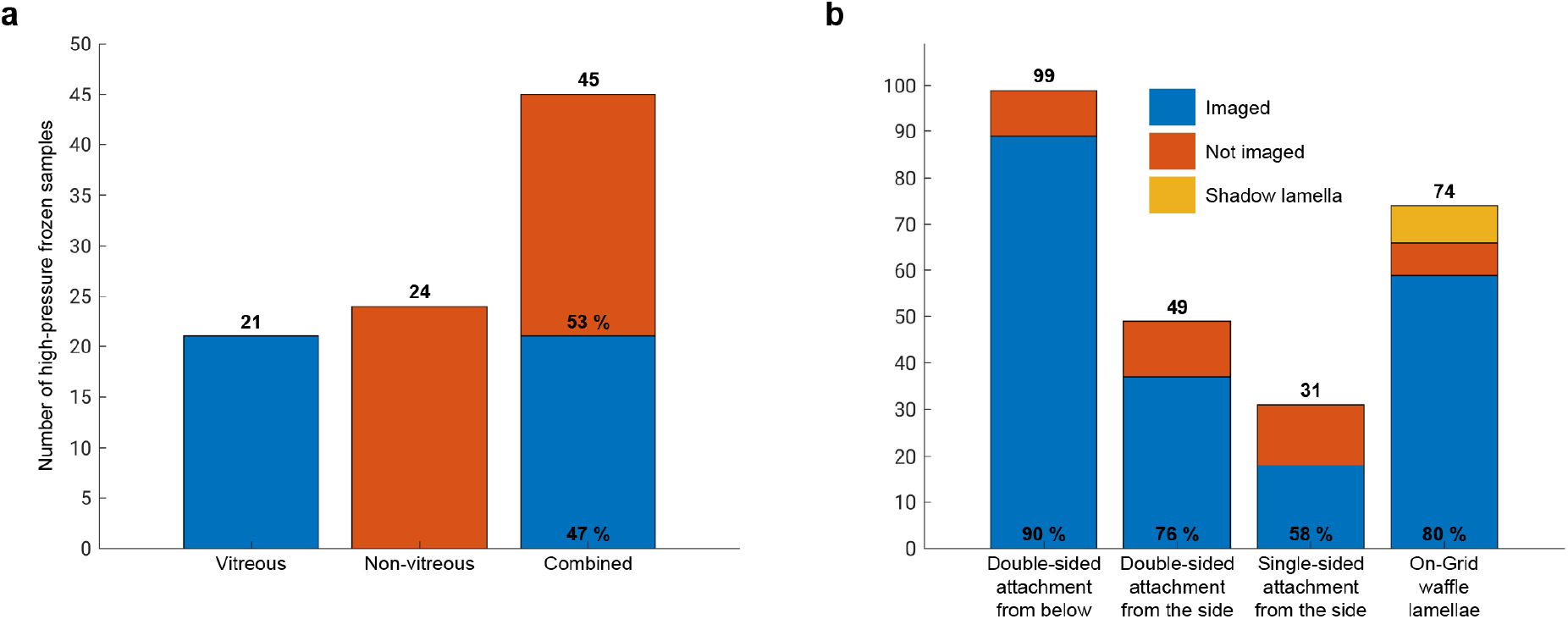
Success rates for high-pressure freezing and lamella preparation. **a**. Success rate for high-pressure freezing for P. patens tissues. **b.** Number of lamellae that were prepared vs the number of imaged lamellae for all tested plant tissues. For on-grid milled waffle lamellae, the fraction “Shadow lamella” indicates lamellae where not all material was removed below the lamella which prevents the acquisition of cryo-ET data.

